# Robinow Syndrome *DVL1* variants disrupt morphogenesis and appendage formation in a Drosophila disease model

**DOI:** 10.1101/2024.09.10.612347

**Authors:** Gamze Akarsu, Katja R. MacCharles, Kenneth Kin Lam Wong, Joy M. Richman, Esther M. Verheyen

## Abstract

Robinow Syndrome is a rare developmental syndrome caused by variants in numerous genes involved in Wnt signaling pathways. We previously showed that expression of patient variants in Drosophila and a chicken model disrupts the balance of canonical and non-canonical/PCP Wnt signaling. We also noted neomorphic effects that warranted further investigation. In this study, we examine morphological changes that occur as a result of one variant, DVL1*^1519^*^Δ*T*^, that serves as a prototype for the other pathogenic variants. We show that epithelial imaginal disc development is disrupted in legs and wings. Shortened leg segments are reminiscent of shortened limb bones seen in RS patients. We find that imaginal disc development is disrupted and accompanied by increased cell death, without changes in cell proliferation. Furthermore, we find altered dynamics of basement membrane components and modulators. Notably we find increased MMP1 expression and tissue distortion, which is dependent on JNK signaling. We also find enhanced collagen IV (Viking) secreted from cells expressing DVL1*^1519^*^Δ*T*^. Through these studies we have gained more insight into developmental consequences of DVL1 variants implicated in autosomal dominant Robinow Syndrome.

## Introduction

Robinow Syndrome (RS) is a rare genetically heterogeneous disorder that is inherited as either an autosomal dominant or recessive disease. The two types can be distinguished by inheritance patterns and severity of phenotypes, with recessive typically leading to more severe clinical outcomes (Mazzeu *et al*., 2007). In dominant RS, pathogenic variants have been identified in 7 genes and all give rise to a set of common phenotypes including “fetal facies” (hypertelorism or wide set eyes, wide nasal bridge, midfacial hypoplasia, micrognathia or smaller mandible and dental irregularities)(Zhang *et al*., 2022). The limbs (especially the forelimbs) are characteristically shorter (mesomelia and brachydactyly) among other skeletal irregularities (Afzal *et al*., 2000; Person *et al*., 2010; Roifman *et al*., 2015; White *et al*., 2015, 2016, 2018; Abu-Ghname *et al*., 2021). The variants primarily affect components of Wnt signaling pathways, including *DVL1*, *DVL2*, *DVL3*, *WNT5A* and *FZD2*. Variants in human Dishevelled 1 (*DVL1*) are the most commonly events implicated in autosomal dominant RS (Zhang *et al*., 2022).

*DVL1* mutations are characterized by a single nucleotide deletion in exon 14 which causes a frameshift that remarkably leads to an extended novel amino acid sequence shared amongst the variants, which replace at least 132 C-terminal amino acids found in wildtype DVL1 (White *et al*., 2015; Hu *et al*., 2022; Lima *et al*., 2022; Zhang *et al*., 2022). We have previously characterized the effects of three such variants (*DVL1^1519^*^Δ*T*^*, DVL1^1529^*^Δ*G*^ and *DVL1^1615^*^Δ*A*^; Fig. 1A) on Wnt signaling pathways using *Drosophila* and chicken models (Gignac *et al*., 2023; Tophkhane *et al*., 2024). By leveraging two model systems we were able to investigate the functional consequences of expression of *DVL1* variants relative to a reference wildtype human *DVL1* gene. We found that the variants disrupted the normal balance found in developing tissues between canonical β-catenin dependent Wnt signaling and non-canonical or planar cell polarity (PCP) Wnt signaling (Gignac *et al*., 2023).

**Fig. 1.**
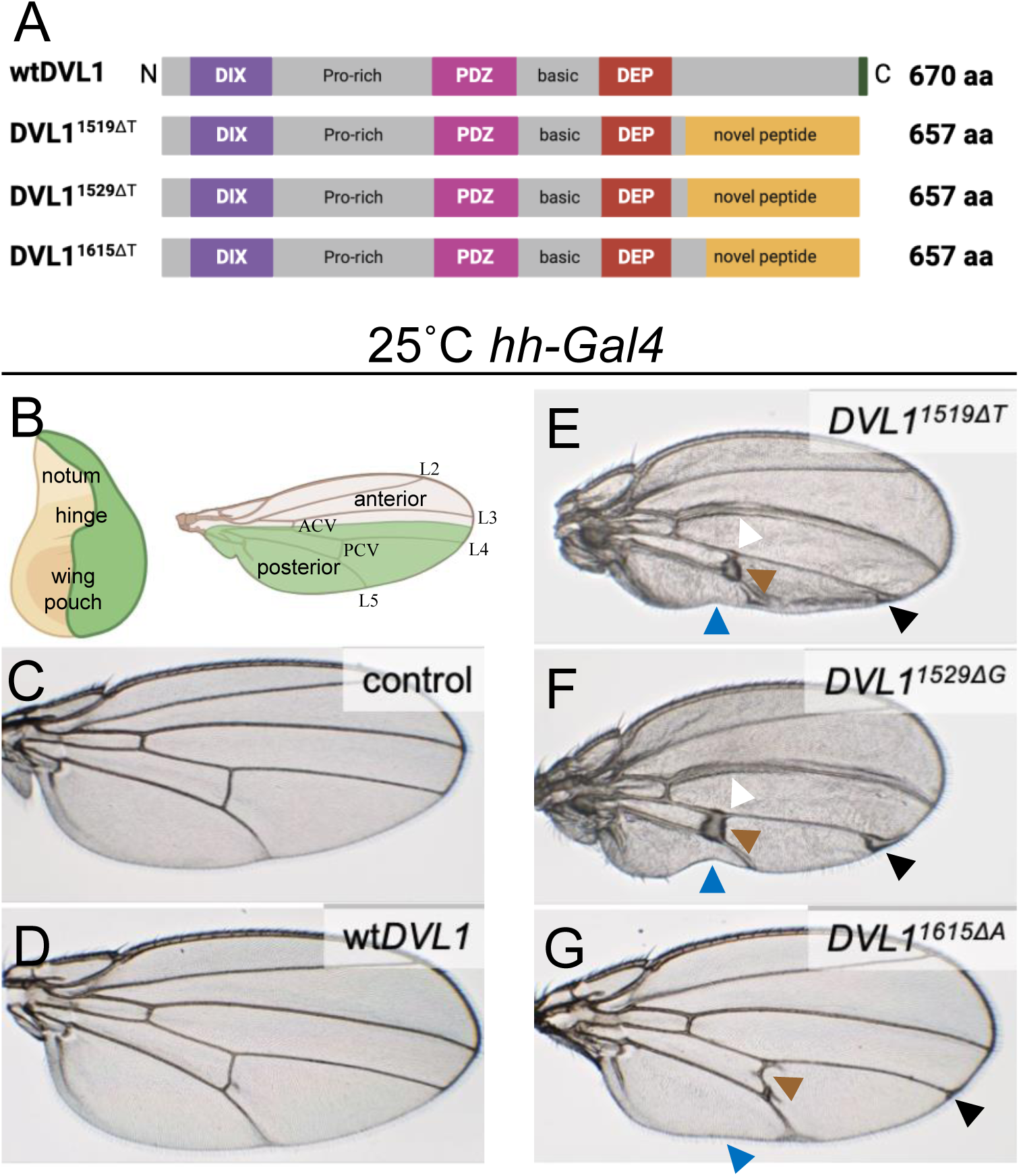
*DVL1^1519ΔT^* serves as a proxy for Robinow Syndrome variants. (A) Graphical representation of human DVL1 protein and three variants associated with Robinow Syndrome. In each variant, a single nucleotide deletion results in a frameshift which causes the addition of a lengthy novel peptide (indicated in yellow) which is shared in large part between all three variants. (B) The *hedgehog-Gal4* (*hh-Gal4*) driver is expressed in the posterior domain of wing imaginal discs which corresponds to the posterior half of adult wings (indicated by green shading). The presumptive notum, hinge and wing pouch domains are indicated in the disc. Longitudinal veins on the wing blade are indicated by L2-5 while the anterior and posterior crossveins are indicated by ACV and PCV, respectively, on the adult wing. (C-G) Representative adult female wings from crosses grown at 25°C of the following genotypes: (C) control *hh>RFP* (D) *hh>wtDVL1*, (E) *hh>DVL1^1519^*^Δ*T*^, (F) *DVL1^1529^*^Δ*G*^ *and* (G) *DVL1^1615^*^Δ*A*^. (E-G) Phenotypes observed were indicated by arrowheads of different colors: longitudinal vein thickening by white, crossvein thickening by brown, distortion in the wing blade shape by blue and vein thickening around the wing margin by black arrowheads.

In our studies in developing *Drosophila* tissues, we expressed the human genes using the Gal4-UAS system which allowed us to precisely control the expression within domains of developing imaginal discs (Brand and Perrimon, 1993). We primarily characterized the effects in a small stripe of cells along the anterior-posterior boundary of the wing imaginal discs using the *dpp-Gal4* driver (Gignac *et al*., 2023). In those studies, we were able to detect decreases in canonical Wnt pathway readouts as well as increased JNK/PCP signaling as assayed by JNK target gene expression as well as detecting the polarity of wing hairs. However, due to the small expression domain, the overall effects on development and viability were mild. We found that ubiquitous expression of DVL1 transgenes causes larval lethality, precluding an analysis of the effects. Thus, there is a need to evaluate the effects on tissue morphogenesis and appendage formation using other tools that allow us to study tissue development without causing early lethality.

In this study we demonstrate that expression of *DVL1* RS mutations in broader domains compared to using *dpp-Gal4* disrupts morphogenesis of appendages and leads to severe disruptions in Drosophila development. Our studies revealed that the *DVL1^1519^*^Δ*T*^ variant can serve as a proxy for the other variants we previously studied and can reveal insights into changes in morphogenesis. Notably, we find severe disruptions in wing and leg formation that reveal that elevated cell death contributes to altered morphology while we found no elevated proliferation. The effect on tissue morphology and expression of MMP1 are dependent on intact JNK signaling. Further, expression of the variant disrupts basement membrane integrity and cell apposition during metamorphosis. These effects provide insight into disruptions that may lead to altered limb formation in RS patients.

## Results

### *DVL1^1519^*^Δ*T*^ serves as a proxy for Robinow Syndrome *DVL1* variants

Our previous work has shown that when *DVL1* variant expression was driven by *dpp- Gal4*, the adult wings displayed planar polarity defects as well as additional mild neomorphic phenotypes such as longitudinal vein thickening, abnormalities in the anterior cross vein, ectopic bristles on the wing margin of the L3 longitudinal vein and a crease between the L3-L4 veins.

These phenotypes were exclusive to the variants only (Gignac *et al*., 2023). Since the *dpp-Gal4* expression domain corresponding to the adult tissue is quite narrow, we wanted to express the variants by using drivers with larger expression domains. We also modulated the temperature at which the flies were raised, since Gal4 has enhanced activity at higher temperatures (Duffy, 2002). We first used *hedgehog-Gal4* (*hh-Gal4*) which is expressed in the posterior compartment of the wing and leg imaginal discs to assess phenotypes caused by human DVL1 proteins at 25°C (Fig. 1B). As shown in our previous work, wt*DVL1* had no effect on wing development (Fig. 1D). In contrast, *DVL1^1519^*^Δ*T*^*, DVL1^1529^*^Δ*G*^ and *DVL1^1615^*^Δ*A*^ all caused disruptions of the posterior domain in adult wings, which included thickening and of longitudinal and cross veins, severe reduction in wing tissue and some distortion of wing blade shape and flatness (Fig. 1E- G). Based on these findings, and our previous work, we chose to focus subsequent studies on characterizing the effects of *DVL1^1519^*^Δ*T*^ as a proxy for the three variants since it consistently phenocopied effects from *DVL1^1529^*^Δ*G*^. We showed that *DVL1^1615^*^Δ*A*^ protein was less stable than the other variants, however when we drive its expression at higher levels, we observed similar effects as those found with the other variants (Gignac *et al*., 2023). Thus, we are focusing here only on *DVL1^1519^*^Δ*T*^ as a proxy for the other pathogenic variants to gain insight into the functional consequences of an RS variant on development.

### *DVL1^1519^***^Δ^***^T^* induces abnormal morphology in the notum and appendages

In examining phenotypes, we scored both females (Fig. 2) and males (Fig. S1). When expressed with the *hh-Gal4* driver at 25°C, the *DVL1^1519^*^Δ*T*^ variant caused the development of smaller, blistered wings that had abnormally enlarged veins (Fig. 1E). In addition to these wing phenotypes, the flies had difficulty eclosing from their pupal cases and often could not walk, leading them to die after falling into the food due to an abnormal leg morphology (Fig. 2C, Fig. S1C). All 3 pairs of legs (termed T1 for pro-, T2 for meso- and T3 for metathoracic leg) were malformed. The most distal part of the Drosophila leg called the tarsus contains 5 tarsal segments. In flies that express the *DVL1^1519^*^Δ*T*^ variant in the *hh-Gal4* domain, the tarsi of T3 often formed a curved structure that appears like a hook (Fig. 2C’, Fig. S1C’) which is not seen in wildtype flies (Fig. 2A’, Fig. S1A’) or those expressing reference wt*DVL1* (Fig. 2B’, Fig. S1B’).

**Fig. 2.**
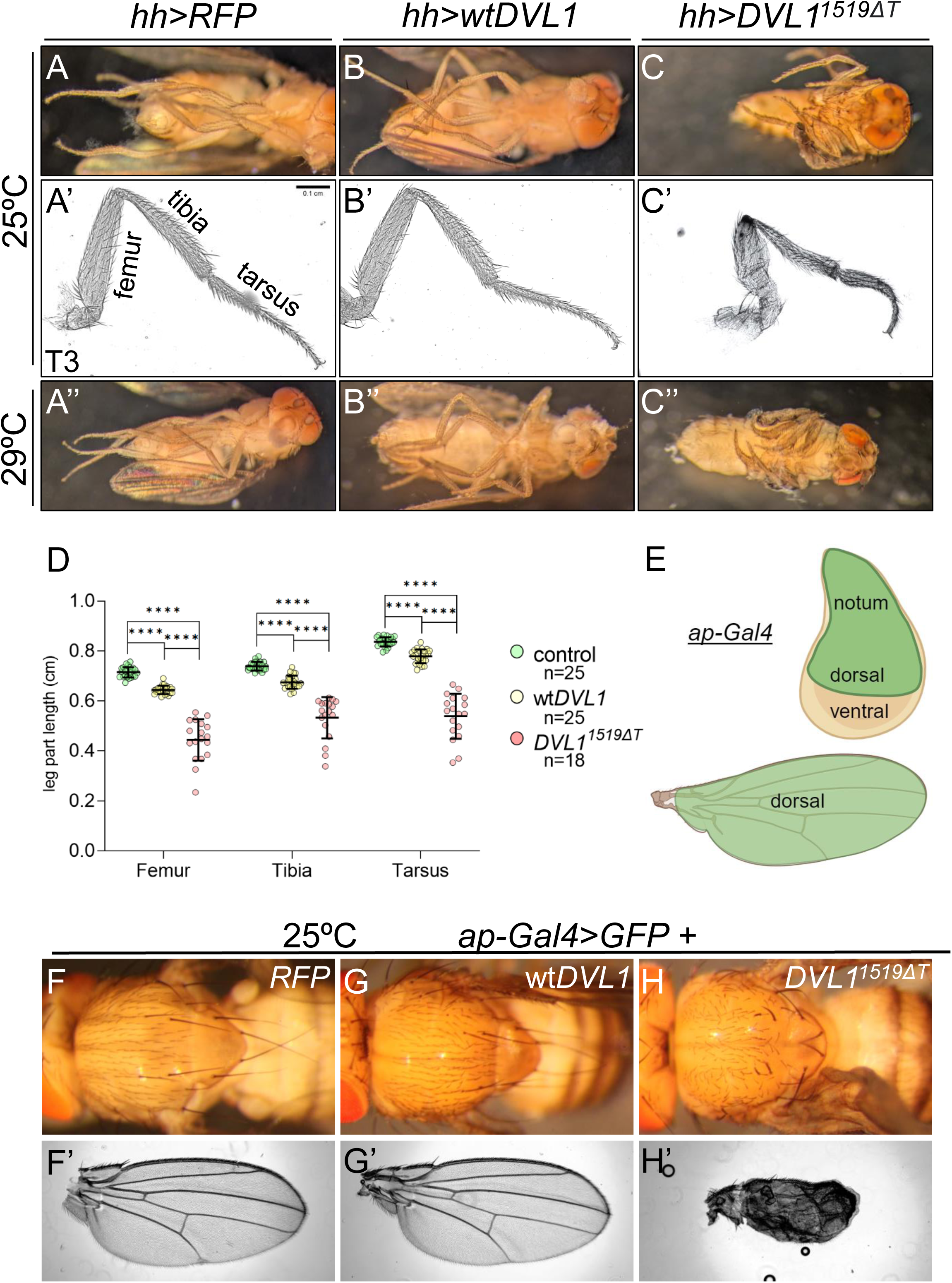
*DVL1^1519^*^Δ*T*^ induces abnormal morphology in female adult fly appendages and thorax. (A-C) Female adults from crosses grown at 25°C of the indicated genotypes. Leg and wing defects are visible in the variant (C). Single third legs (T3) showing of the indicated genotypes. (A”-C”) Females from crosses grown at 29°C to drive higher transgene expression. Pharate pupae in C” was dissected from pupal case. (D) The length of T3 leg segments were measured and plotted for control, wt*DVL1* and *DVL1^1519^*^Δ*T*^ expressing flies, indicated by green, yellow and pink dots, respectively. Statistics were performed using Tukey’s multiple comparisons test. ****p<0.0001. (E) The *apterous-Gal4* (*ap-Gal4*) driver is expressed in the dorsal domain of wing imaginal discs which corresponds to the dorsal side of adult wing blades (indicated by green shading). The presumptive notum, hinge, wing pouch domains are indicated in the disc. (F-H) Female adult body parts from crosses of the indicated genotypes grown at 25°C. (F-H) The adult notum and scutellum. (F’-H’) Adult wings of indicated genotypes.

Furthermore, the femur and tibia segments were also shorter and appeared to have disrupted planar polarity (Fig. 2C’, Fig. S1C’). When measured, we found that the length of all three leg segments were significantly shorter in *DVL1^1519^*^Δ*T*^ expressing flies compared to both control and wt*DVL1* expressing flies (Fig. 2D, Fig. S1D).

When we expressed *DVL1^1519^*^Δ*T*^ with *hh-Gal4* at higher levels by growing the flies at 29°C, we observed more severe phenotypes compared to the flies grown at 25°C (Fig. 2C”, Fig. S1C’’). Variant expressing flies died inside their pupal cases at the pharate adult stage. Since the expansion of wings occurs after eclosion, we were not able to image the adult wings. Variant expressing pupae were dissected out of their cases to document the disrupted morphogenesis (Fig. 2C”, Fig. S1C’’). None of the wt*DVL1* expressing flies had difficulties eclosing at 29°C nor showed any obvious phenotypes (Fig. 2B”, Fig. S1B’’).

We next expressed the *DVL1^1519^*^Δ*T*^ variant with the *ap-Gal4* driver which is expressed in the dorsal compartment of the wing imaginal disc which gives rise to one side of the wing blade and the adult notum (or thorax) (Fig. 2E). During morphogenesis the two wing discs normally fuse to generate a seamless notum with a small posterior scutellum structure. Expression of the *DVL1^1519^*^Δ*T*^ variant disrupted wing development and wing expansion did not occur properly after eclosion, leading to shrivelled and blistered wings (Fig. 2H’). The thorax of the flies was also abnormally developed (Fig. 2H, Fig. S1G-G’’). The flies had missing or malformed scutella and some of the thorax bristles were missing, and the remaining bristles showed PCP defects. No leg malformations were observed since *ap-Gal4* is not expressed in leg discs, enabling flies to eclose.

### *DVL1^1519^***^Δ^***^T^* disrupts morphology in larval wing imaginal discs

The phenotypes observed in adult flies with the expression of the *DVL1^1519^*^Δ*T*^ variant led us to investigate the morphology of wing precursor tissues in larvae. We investigated the wing imaginal discs that expresses the variant using *ap-Gal4* and *hh-Gal4* drivers. We used *UAS-GFP* in these crosses to mark the domains in which transgenes were expressed. Imaginal discs comprise a columnar epithelium (disc proper) and a thin squamous peripodial cell layer. In our studies we focussed solely on the disc proper.

*hh-Gal4* was used to drive expression of a control *UAS-RFP* line, wt*DVL1* and *DVL1^1519^*^Δ*T*^ at 29°C and imaginal discs were dissected from male and female third instar larvae and stained with DAPI and for F-actin to reveal tissue morphology (Fig. 3A-C). Orthogonal views of discs reveal the apical-basal orientation of the disc proper (small insets in 3A’-C’). Expression of wt*DVL1* did not disrupt wing disc morphology (Fig. 3B-B’) and orthogonal views showed a normal disc structure. The activity of *DVL1^1519^*^Δ*T*^ in the posterior side of the wing imaginal discs induced extra folds in every direction, making the wing disc appear highly distorted (Fig. 3C). All parts of the wing disc, notum, hinge and wing pouch showed abnormal morphology. The orthogonal view from showed disruption of the arc-like form of the wing discs. With all the extra folds, the wing discs appeared highly overgrown and multilayered. The tissue in the presumptive notum and hinge above the wing pouch (labeled in Fig. 1B) appeared highly misfolded on the basal surface of the wing imaginal discs (Fig. 3C’). These phenotypes were also observed when the *DVL1^1519^*^Δ*T*^ expression was lower at 25°C (Fig. 2S) but they were more striking at 29°C.

**Fig. 3.**
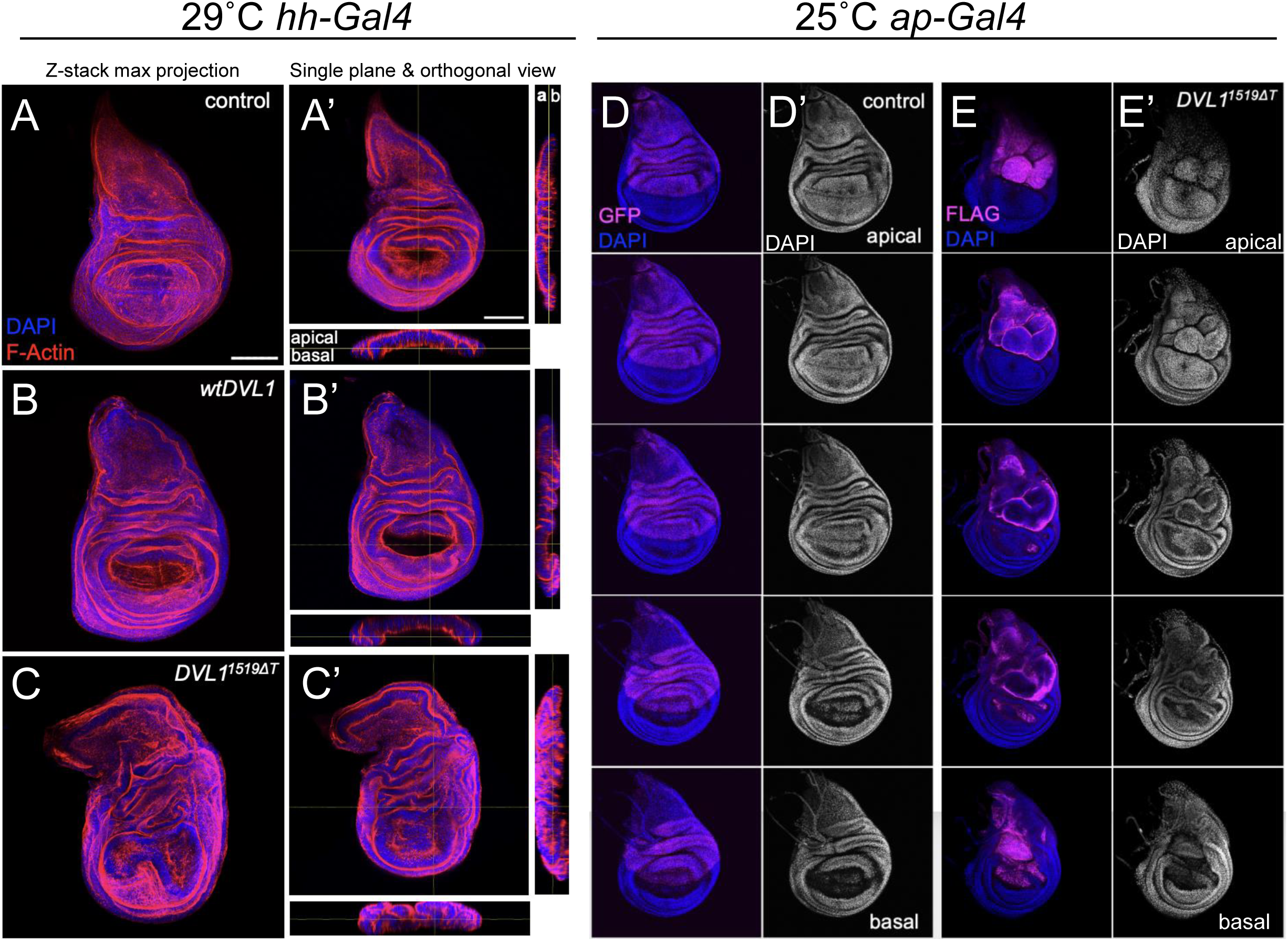
*DVL1^1519^*^Δ*T*^ variant disrupt morphology in larval wing imaginal discs. (A-C) Wing imaginal discs from female larvae grown at 29°C and using *hh-Gal4* to express transgenes. Images showing DNA (DAPI, blue), F-actin (red) for (A, A’) control *UAS-RFP* (B-B’) wt*DVL1* and (C-C’) *DVL1^1519^*^Δ*T*^ wing discs mounted on bridged slides to retain their three- dimensional shape. (A-C) Maximum projections, (A’-C’) single plane images with orthogonal views of the tissue. Scale bar = 100 μm. (D-E) Wing imaginal discs from female larvae grown at 25°C and using *ap-Gal4* to express control and *DVL1^1519^*^Δ*T*^ transgenes, as indicated. DAPI (blue) and anti-FLAG or GFP (magenta) were used to show the cells expressing transgenes in merged images (D, E) while single channel DAPI (white) staining shown is in D’, E’ to highlight tissue morphology. Single plane images are organized apical to basal as labelled.

When the *DVL1^1519^*^Δ*T*^ variant was expressed in the dorsal compartment with *ap-Gal4* at 25°C, we observed that the expression domain in the wing disc, shown with the FLAG antibody staining against the N-terminal FLAG-tag of DVL1 proteins expressed, was distorted, and the dorso-ventral border was expanded towards the ventral compartment (Fig. 3E-E’) compared to control discs (Fig. 3D-D’). By showing single sections in a progression from apical to basal we can reveal the distortion in the wing tissue (Fig. 3D’, E’). We also observed that the highly regulated folding structure of the wing imaginal disc was altered, and more layers appeared to be formed above the wing pouch especially on the posterior side of the wing imaginal discs. The abnormal folding was also present on the dorsal domain, in the notum. These phenotypes, especially the misfolding, were also observed when *DVL1^1519^*^Δ*T*^ was expressed in higher levels at 29°C (Fig. 3S).

### *DVL1^1519^*^Δ*T*^ induces apically and basally shifted cells

In Fig. 3, we showed that the tissue was distorted along the apico-basal axis. To investigate this in more detail we generated RFP-marked flip-out clones expressing either control (*white-RNAi*, Fig. 4A-B), wt*DVL1* (Fig. 4C-D) or *DVL1^1519^*^Δ*T*^ (Fig. 4E-F) using *actin-Gal4* in wing discs. Control and wt*DVL1* clones both showed normal tissue architecture, apico-basal polarity, E-cadherin protein levels and localization. In contrast, *DVL1^1519^*^Δ*T*^-expressing clones were apically and basally shifted (Fig. 4F”). In some cases, cells appear to be extruding from the basal side of the disc. In addition to the decreased levels of β-catenin/Armadillo which we previously found after expressing *DVL1^1519^*^Δ*T*^ (Gignac *et al*., 2023), we now show that *DVL1^1519^*^Δ*T*^ expression was accompanied by a mild decrease in E-cadherin (E-cad) staining (Fig. 4F’) and reduced E-cad at apical junctions (Fig. 4F”). Considering β-catenin/Armadillo’s essential role in the adherens junction assembly (Cox *et al*., 1996), the decrease in E-cadherin levels is in line with our earlier findings.

**Fig. 4.**
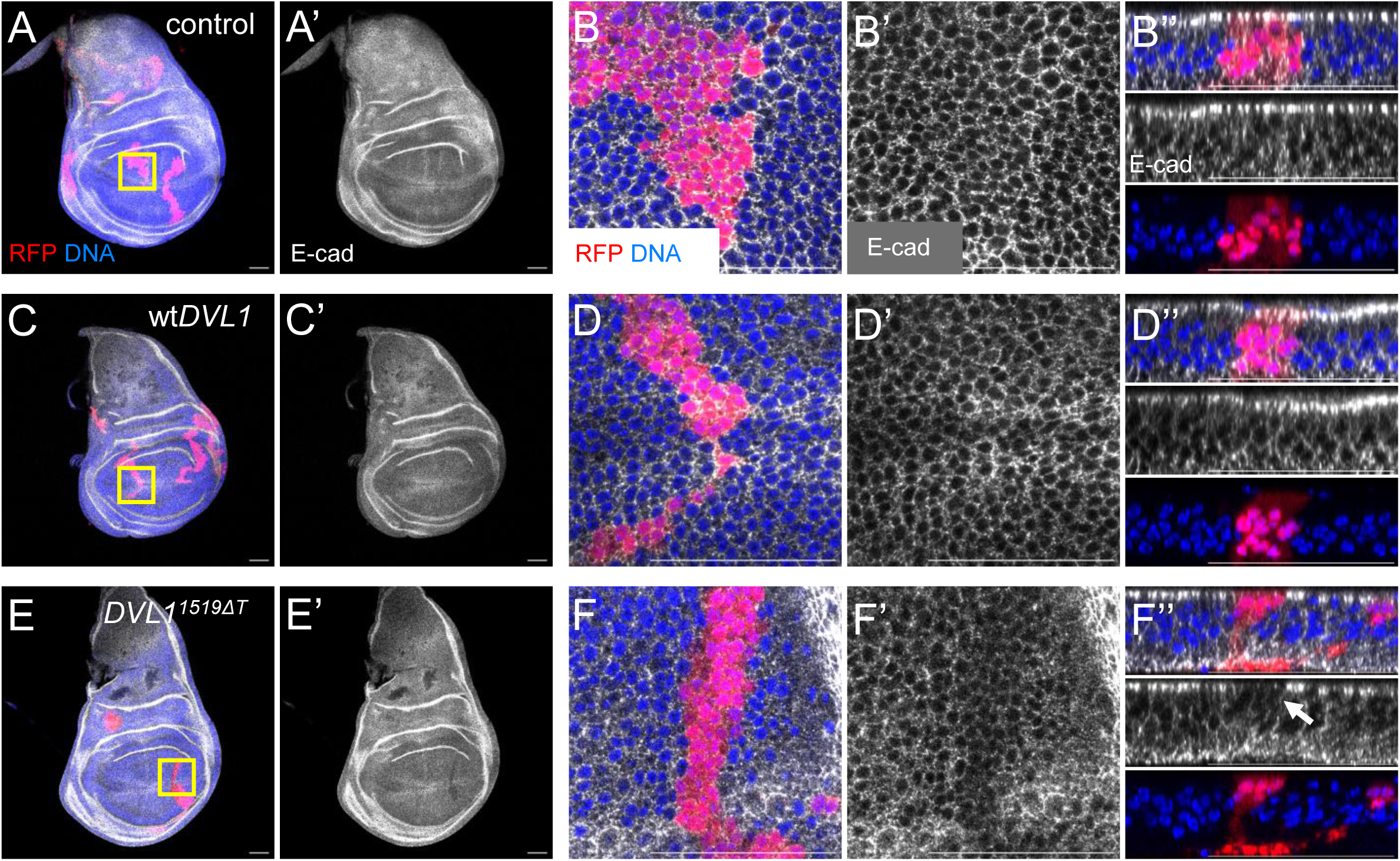
Epithelial organization is disrupted by *DVL1^1519^*^Δ*T*^. (A-B”) Control heat shock inducible flip out clones in wing discs are positively marked with RFP. DNA is labelled with DAPI (blue). (B, D, F) Zoomed in regions highlighted by yellow boxes in A, C, E. (B-B”) E-cadherin protein localization in at the apical surface (B’) and orthogonal view (B”) shows apical enrichment in columnar epithelial cells of disc. Cells in flip out clones are marked with RFP and extend along the apical-basal axis of the disc. (C-D”) Clones of cells expressing wt*DVL1*. (F-F”) Clones of cells expressing *DVL1^1519^*^Δ*T*^. In F”, cells are seen at the apical and basal surfaces of disc, and E-cadherin staining appears reduced (arrow). Scale bar = 50 μm.

### *DVL1^1519^***^Δ^***^T^* does not induce cell proliferation in wing imaginal discs

After observing the highly altered morphology in wing imaginal discs that express the *DVL1^1519^*^Δ*T*^ variant in *apterous* and *hedgehog* domains, we wanted to investigate whether cell proliferation was being induced to cause the extra fold formation. To assess the relative amount of cell proliferation occurring between the wt*DVL1* and variant expressing wing imaginal discs, we used an antibody against a commonly used specific mitotic marker, Phospho-histone H3 (PH3).

We investigated the levels of cell proliferation in wing imaginal discs that expresses either wt*DVL1* or *DVL1^1519^*^Δ*T*^ variant in the *hedgehog* domain in comparison to the control discs at 25°C (Fig. 5A-C”). The number of PH3 puncta in all three genotypes were very similar and the variance was very small (Fig. 5D). Statistical analyses showed no significant difference between the discs that express the *DVL1^1519^*^Δ*T*^ or wt*DVL1* and the control. From these results, we concluded that the abnormal morphology of the wing imaginal discs induced by the expression of *DVL1^1519^*^Δ*T*^ variant is not due to cell proliferation or tissue overgrowth. These results were confirmed by experiments conducted by expressing the *DVL1^1519^*^Δ*T*^ variant in the *hedgehog* domain at 29°C and in the apterous domain at both 25°C and at 29°C (Fig. S4, Fig. S5, Fig. S6, respectively).

**Fig. 5.**
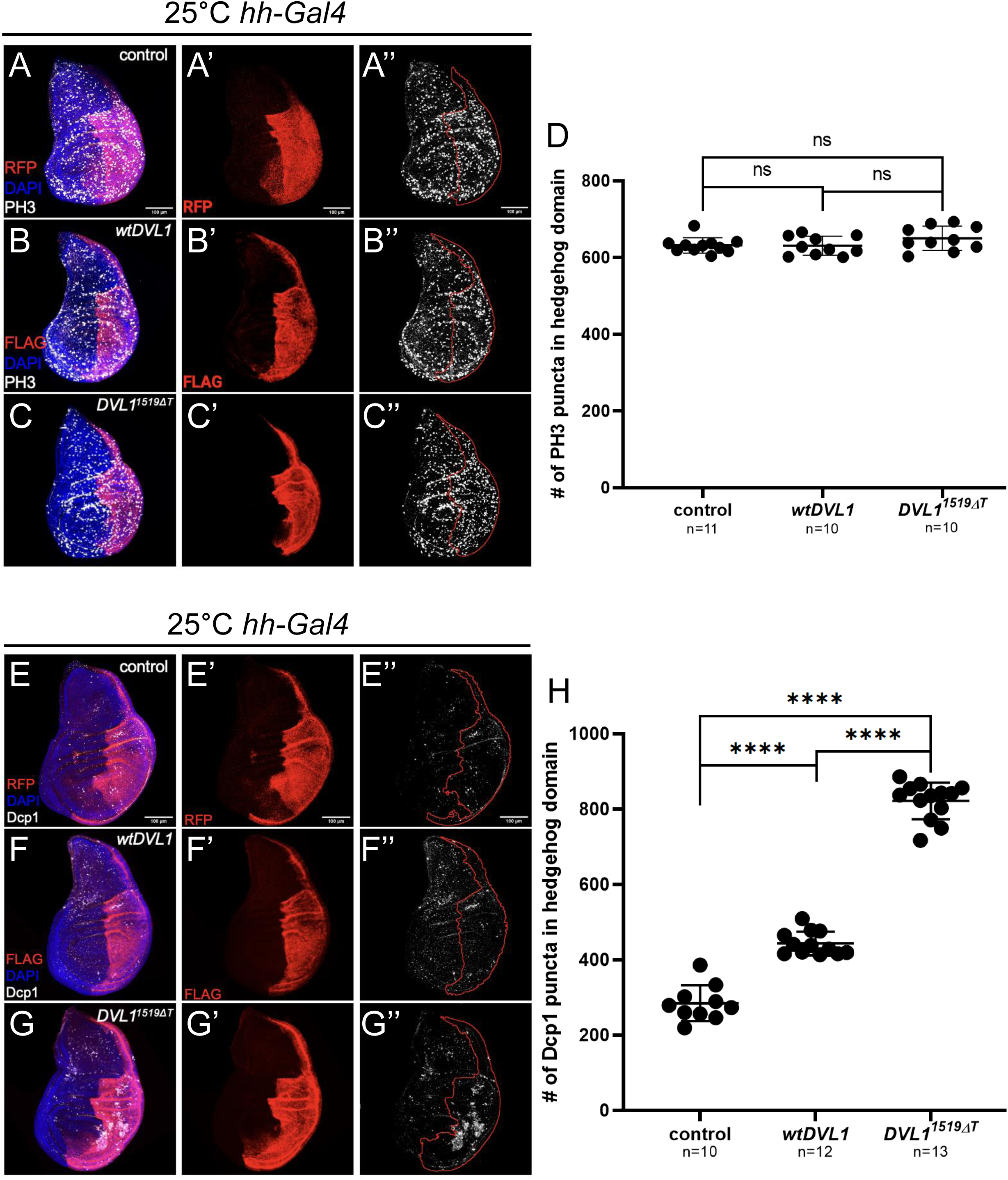
*DVL1^1519^*^Δ*T*^ does not promote proliferation but increases cell death. (A-C) Merged confocal microscopy images showing the *hh-Gal4* UAS transgene expression domain (anti-FLAG or RFP, red), DNA (DAPI, blue) and PH3 (white) staining in (A) control, (B) wt*DVL1* and (C) *DVL1^1519^*^Δ*T*^ wing discs grown at 25°C. (A’, B’, C’) The *hh-Gal4* domain of representative wing discs is shown with the RFP or FLAG staining images. (A’’, B”, C’’) PH3 staining for all genotypes are shown. (D) The number of PH3 puncta counted within the *hh-Gal4* domain are plotted. (E-G) Merged confocal microscopy images showing the *hh-Gal4* UAS transgene expression domain (anti-FLAG or RFP, red), DNA (DAPI, blue) and anti-Dcp1 (white) staining in (E) control, (F) wt*DVL1* and (G) *DVL1^1519^*^Δ*T*^ wing discs grown at 25°C. (H) The number of Dcp1 puncta counted within the *hh-Gal4* domain are plotted. Sample size is shown under each genotype. Statistics were performed with ANOVA. ****p<0.0001. Scale bar = 100 μm.

### *DVL1^1519^*^Δ*T*^ induces cell death in wing imaginal discs

To further investigate the reasons underlying the altered morphology in the wing imaginal discs and the adult tissues induced by the expression DVL1 variant, we investigated the relative amount of cell death occurring between wild type and variant forms of *DVL1*. Our earlier work has found that the DVL1 variant expression causes an imbalance in Wnt signalling and increases the activity of noncanonical Wnt signalling, which mediates cytoskeletal organization and apoptosis (Gignac *et al*., 2023). We also showed that when DVL1 variants were expressed, JNK signalling was ectopically induced, which also regulates apoptosis (Gignac *et al*., 2023). Considering these alterations in signalling and the smaller thorax and adult wing phenotypes induced by the expression of DVL1 variant in different domains, we scored cell death in larval wing imaginal discs using an antibody against the cleaved Death caspase-1 (Dcp1) protein (Song, McCall and Steller, 1997), which is an effector caspase, activated after the cell goes into apoptosis.

We induced the expression of either RFP, wt*DVL1* or *DVL1^1519^*^Δ*T*^ by using the *hh-Gal4* driver at 25°C (Fig. 5E-G”). We counted the number of Dcp1-expressing puncta in the posterior domain of the wing imaginal discs (Fig. 5H). We found that both wt*DVL1* and *DVL1^1519^*^Δ*T*^ expression caused a significant increase in the Dcp1 puncta numbers compared to the control discs (Fig. 5H). *DVL1^1519^*^Δ*T*^ variant expressing discs had significantly higher numbers of Dcp1 puncta than the wt*DVL1* expressing discs (Fig. 5H).

With these results, we conclude that the expression of the *DVL1* RS variant leads to increased levels of apoptosis in wing imaginal disc epithelia. These results support our finding that the abnormal morphology is not caused due to an increase in cell proliferation. *DVL1^1519^*^Δ*T*^ expression also disrupts junctional protein organization as we showed for E-cadherin (Fig. 4F’- F’’).

### *DVL1^1519^*^Δ*T*^ effects depend on JNK signaling

To probe if *DVL1^1519^*^Δ*T*^-induced phenotypes are caused by JNK activation downstream of PCP signaling, we expressed the dominant negative form of Basket (*Drosophila* JNK; bsk^DN^) (Adachi-Yamada *et al*., 1999) to inhibit JNK signaling while expressing the RS variant. We then monitored two well established targets of JNK signaling, the transcriptional reporter *puckered-lacZ* (*puc-lacZ)* (Martin-Blanco *et al*., 1998) and Mmp1 protein levels (Uhlirova and Bohmann, 2006) (Fig. 6). We first used *ap-Gal4* to drive expression of *DVL1^1519^*^Δ*T*^ and then co-expressed either a benign *UAS-GFP* (Fig. 6A-B) or the *UAS-bsk^DN^* transgene (Fig. 6C-D). Wing discs were stained with DAPI to reveal overall tissue morphology, and the expression of *puc-LacZ* and the *DVL1* transgene were detected with antibodies against β-galactosidase (Fig. 6A’-D’) and the FLAG tag (Fig. 6A”-D”), respectively. Expression of *DVL1^1519^*^Δ*T*^ and GFP revealed the ectopic *puc-lacZ* staining and distorted *ap-Gal4* domain that we previously reported (arrow in Fig. 6B, Fig. 6B’)(Gignac *et al*., 2023). In contrast, co-expressing *bsk^DN^* with the *DVL1* variant suppressed the ectopic *puc-lacZ* expression and suppressed tissue distortion (Fig. 6C’, arrow in D, D’). We measured the *puc-lacZ* expression levels within and outside the *DVL1^1519^*^Δ*T*^ expression domain in the wing discs and plotted the fold change of signal intensity which showed that the co-expression of *bsk^DN^* significantly decreased the ectopic *puc-lacZ* expression levels (Fig. 6I).

**Fig. 6.**
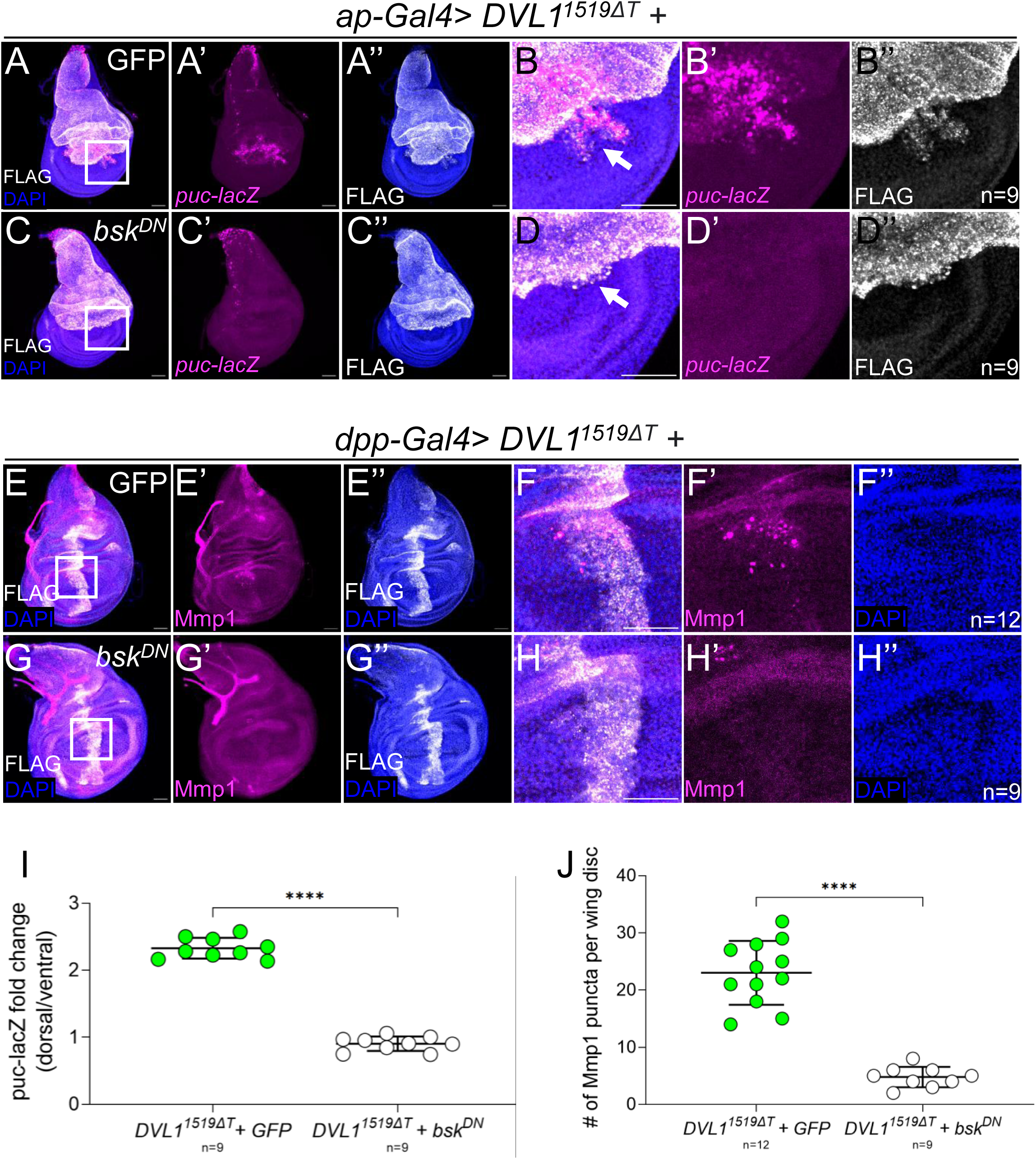
DVL1^1519ΔT^ effects are mediated through JNK signaling. (A-D’) *ap-Gal4* was used to express *DVL1^1519^*^Δ*T*^along with either control *UAS-GFP* (A-B”) or *UAS-bsk^DN^* (C-D”). (B, D) Magnified views of regions shown in white boxes in A and C. Wing discs were stained with DAPI to reveal tissue organization, and the transcriptional reporter *puckered-lacZ* (*puc-lacZ*) as a readout of Jnk signalling. Anti-FLAG antibodies were used to reveal cells expressing transgenes. (E-H”) dpp-Gal4 was used to express *DVL1^1519^*^Δ*T*^along with either control *UAS-GFP* (E-F”) or *UAS-bsk^DN^* (G-H”). (F, H) Magnified views of regions shown in white boxes in E and G. Antibodies were used to detect Mmp1 protein levels and FLAG. (I) The fold change of puc-lacZ signal intensity inside and outside the domain expressing *DVL1^1519^*^Δ*T*^ (shown in A-D’) were measured and plotted for both genotypes. (J) Mmp1 puncta per wing disc were counted and plotted. Statistics were performed using Welch’s t-test. ****p<0.0001. Scale bar is 100 μm for E-E’’, G-G’’; 20 μm for F-F’’, H-H’’.

We carried out a similar experiment using *dpp-Gal4* to drive *DVL1^1519^*^Δ*T*^ in a narrow stripe along the anterior posterior boundary of the wing disc (Fig. 6E-H). We previously showed that expressing any of the three DVL1 variants with *dpp-Gal4* leads to ectopic Mmp1 expression (Gignac *et al*., 2023), which we show here with *DVL1^1519^*^Δ*T*^ (Fig. 6E’, F’). This effect was reversed by co-expression of *bsk^DN^* (Fig. 6G’, H’). We counted the numbers of ectopic Mmp1 puncta in the wing discs expressing *DVL1^1519^*^Δ*T*^ in the *dpp* domain and plotted the results.

Statistical analysis showed that inhibiting JNK signaling caused a significant decrease in the Mmp1 puncta present in the wing discs (Fig. 6J). The results from these two experiments shows that the RS variant effects on tissue morphology and ectopic induction of *puc-lacZ* and Mmp1 are dependent on JNK signaling.

### *DVL1^1519^*^Δ*T*^ disrupts cell adhesion in pupal wings

We found that *DVL1^1519^*^Δ*T*^ caused dramatic phenotypes in adult wings by altering wing development, creating enlarged veins and decreasing the overall size of the wing blade. We also observed numerous defects in tissue morphology in larval imaginal discs. Based on these results, we wanted to examine pupal wings to gain insight into alterations happening during metamorphosis. Pupal wing developmental stages are well described for flies grown at 25°C (Waddington, 1940; Diaz de la Loza and Thompson, 2017; Bier, 2000). We examined the wings of pupae 28-30 hours after puparium formation (APF) since the wings at this stage have actin hairs and veins and reflect the “definitive shape” of the adult wings.

We used *dpp-Gal4* to express the control, wt*DVL1* and the RS variant along with UAS-GFP in a narrow domain which corresponds to the region between the presumptive 3^rd^ and 4^th^ longitudinal veins (L3 and L4; Fig. 7A). Expression of wtDVL1 had no effect on pupal wing morphology as revealed by staining for F-actin (Fig. 7B-B’). When *DVL1^1519^*^Δ*T*^ was expressed in this domain, it caused an apparent blistering phenotype in the 28-30 hr APF pupal wings (Fig. 7C-C’, arrow). At this stage of the pupal wing development, the vein pattern is established. This blister phenotype appeared to be due to a failure of the dorsal and ventral layers of the wing to adhere together to form the proper vein and wing blade structure. To examine this idea, we looked at a cross section of the wing blade in two regions – one where no DVL1 transgene was expressed (Fig. 7D, box E) and one within the region that appeared to be blistered (Fig. 7D, box E’). In normal wing tissue at 28-30 hrs APF the two sheets of epithelial cells can be seen to be tightly apposed at their basal surfaces, as their cell shapes are outlined by F-actin and DAPI reveals nuclear position (Fig. 7E). In contrast, in the region of the pupal wing that showed a blister, there is a large gap between the two sheets of cells and loose cells can be seen floating in the lumen (Fig. 7E’).

**Fig. 7.**
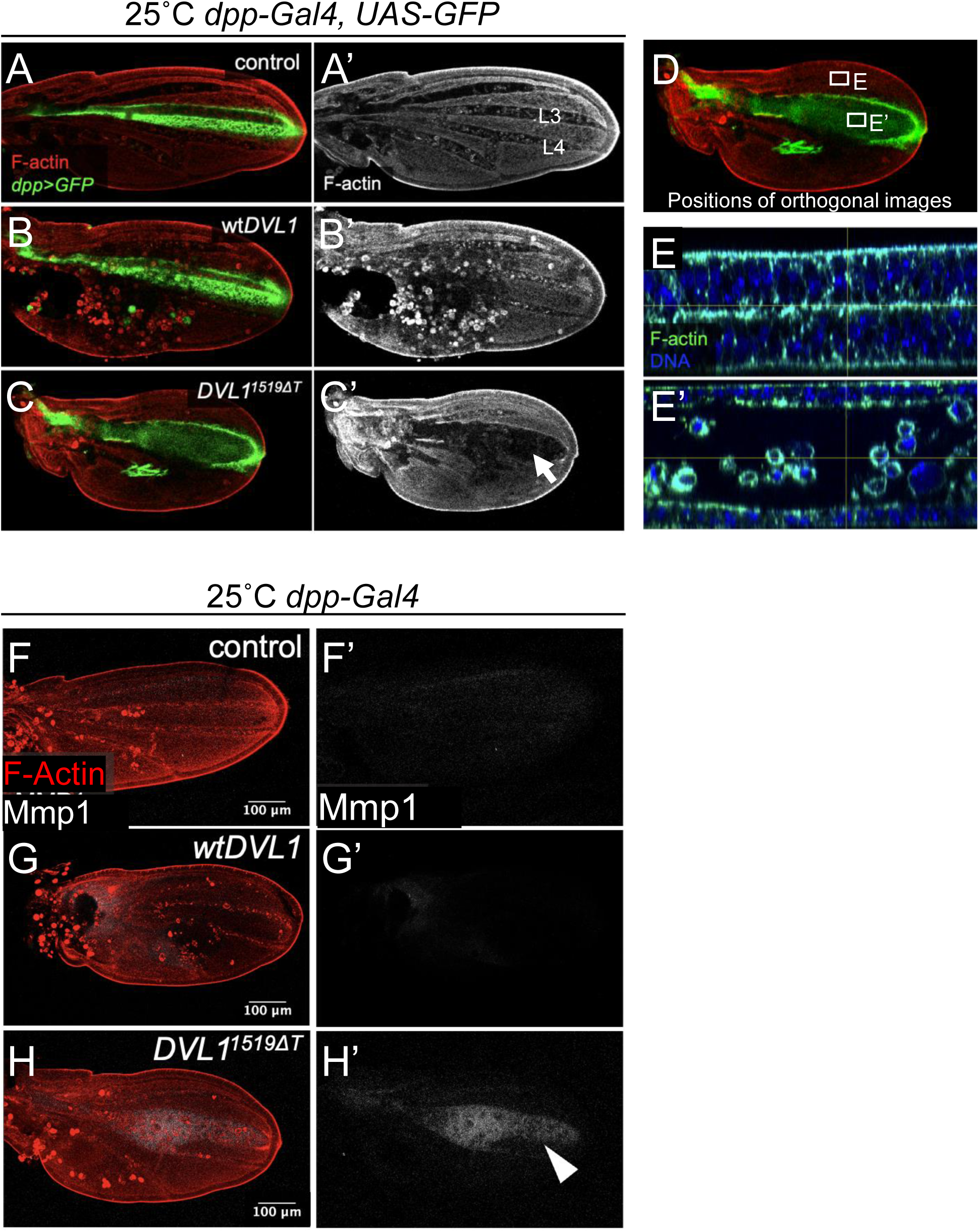
Pupal wing development is disrupted by expression of a DVL1 variant. The *dpp-Gal4>UAS-GFP* expression domain is shown in green and actin (rhodamine-phalloidin, red) staining in (A) control, (B) wt*DVL1* and (C) *DVL1^1519^*^Δ*T*^ pupal wings dissected 28hrs after puparium formation (APF). Single channel actin (white) stain shown in (A’, B’, C’). Arrow indicates pupal wing blister. (D) *DVL1^1519^*^Δ*T*^ pupal wing in which areas shown in E and E’ are indicated. Cross sections outside (E) and inside (E’) of the *dpp-Gal4* expression domain are shown. (E-E’) Single plane images of orthogonal views showing F-Actin (green) and DNA (blue) staining in 28hr APF *dpp>DVL1^1519^*^Δ*T*^ pupal wings above (A) and inside (B) of a wing blister. Crosses were performed at 25°C and female and wings are shown.

An earlier study has shown that the overexpression of Matrix-metalloproteases (Mmps) can induce blister formation in pupal wings (Sun *et al*., 2021). We previously showed that the RS variants induce ectopic Mmp1 expression in wing imaginal discs (Gignac *et al*., 2023). Thus, we investigated if the ectopic expression of Mmp1 is also present in the pupal wing blisters.

Staining of either control or *dpp>*wt*DVL1* pupal wings using antibodies against Mmp1 showed very low-level expression (Fig. 7F’-G’). In contrast, we found that *DVL1^1519^*^Δ*T*^-induced blister shows elevated Mmp1 protein in 28-30hr APF pupal wings (Fig. 7H-H’).

We next looked at the blisters more closely. The proper adhesion of the dorsal and ventral layers of the wings occurs with the slow degradation of the basement membrane (Sun *et al*., 2021). Since we hypothesized that there was an alteration in the adhesion process, we wanted to look at the basement membrane. We chose to target *Drosophila* Collagen IV called *Viking* (*Vkg*) as a proxy for the basement membrane since it is the most abundant component. We used Vkg-GFP expressing flies, which express an endogenously tagged Vkg-GFP fusion protein to visualize the basement membrane in pupal wings.

The variant and wt*DVL1* expression were driven by *dpp-Gal4* driver (Fig. 8B-C). We found that the Vkg-GFP is not abundant in the control pupal wing, we only observed a faint GFP signal withing the veins (Fig. Fig. 8A’-A”). We observed a sharper Vkg-GFP signal in the veins of the wt*DVL1* expressing pupal wings (Fig. 8B’-B”). The vein pattern was mostly comparable to control wings, however, L4 longitudinal vein of wt*DVL1* expressing pupal wings appeared malformed and sometimes bent. The pupal wings that express the *DVL1^1519^*^Δ*T*^ variant had the most intense Vkg-GFP signal (Fig. 8C’-C”). The veins and the area around the veins seemed to be filled with Vkg-GFP. The L3 longitudinal vein is enlarged, forming a blister-like structure. The region between the L3 and L4 longitudinal veins appeared much narrow supporting the idea that

**Fig. 8.**
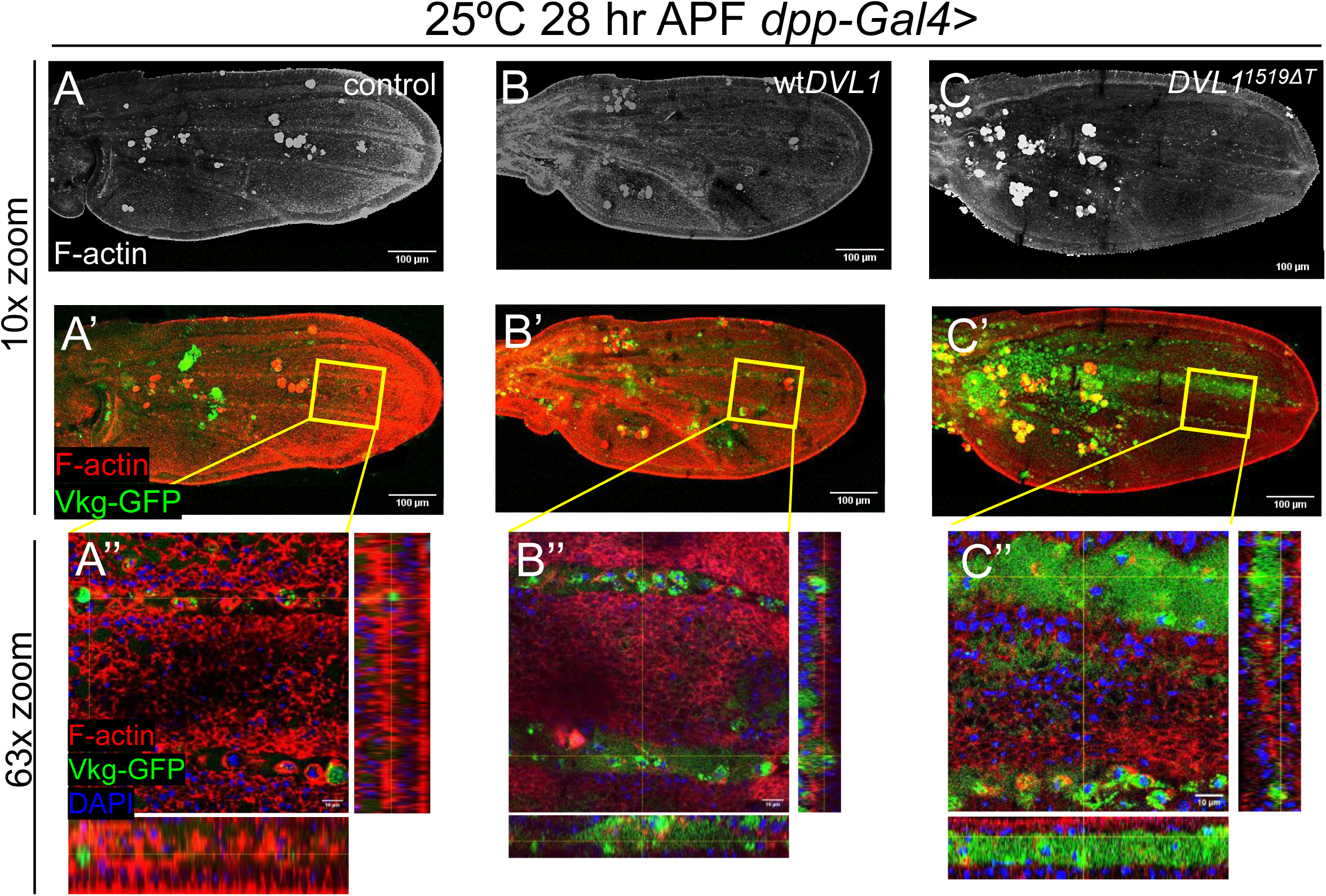
DVL1^1519ΔT^ induces Viking accumulation. 28hr APF pupal wings showing F-Actin (red) and Vkg-GFP (green) staining in control (A), wt*DVL1*- expressing (B) or *DVL1^1519^*^Δ*T*^-expressing (C) tissue at 10x zoom. 63x zoomed in images of the region marked by the yellow squares in A-C are shown in A’-C’ respectively also with DAPI staining showing the cell nuclei in blue. Crosses were performed at 25°C. Scale bar is 100 μm for A-C’; 10 μm for A’’-C’’.

## Discussion

Robinow Syndrome mutations in the *DVL1* gene cause a range of clinical phenotypes including bone malformations (Afzal *et al*., 2000; Person *et al*., 2010; Roifman *et al*., 2015; White *et al*., 2015, 2016, 2018; Abu-Ghname *et al*., 2021). In our previous work we described that *DVL1* frameshift mutations induce an imbalance in Wnt signaling pathways in chick and *Drosophila* models of RS (Gignac *et al*., 2023). In that study we also described several neomorphic phenotypes in adult wings that suggested the variants could disrupt other developmental processes. Among these was an unusual crease in the region between longitudinal veins 3 and 4, as well as disruption in the anterior cross vein. In this current study we have used Drosophila limbs and wings to investigate cellular changes that occur following expression of the prototypical RS variant *DVL1^1519^*^Δ*T*^.

### DVL1 variants induce neomorphic phenotypes

Expression of *DVL1^1519^*^Δ*T*^ in broad domains within developing imaginal discs disrupted development of the organization of the columnar epithelial cells. These effects lead to dramatic effects in leg development, causing shortening of limb segments analogous to bone shortening seen in RS patients. Furthermore, we found defects in the notum that suggested failure to form a seamless scar after the wing imaginal discs fuse during morphogenesis. Wing formation is also severely disrupted, affecting the overall size and shape of the wing blade and structure of the veins. We also observed disruption of planar cell polarity manifested by the bristles on these body parts that were malformed.

*Drosophila* appendage development is governed by well characterized signaling pathways. The adult thorax mostly develops from the wing disc notum, influenced by the balance of Wg and EGFR signaling (Wang *et al*., 2000; Tripathi and Irvine, 2022). Leg formation from embryo to adult is regulated by Dpp, Notch, and Hh signaling pathways (Lecuit and Cohen, 1997; Córdoba *et al*., 2016; Heingård *et al*., 2019), while cross vein patterning in wings depends on BMP and Dpp signaling (Montanari *et al*., 2022). Notably, multiple pathways, including Notch, are involved in bristle shaping and directionality (Furman and Bukharina, 2011). All these literature suggest that the expression of *DVL1^1519^*^Δ*T*^ variant alters the regular functioning of signaling pathways. The mechanisms in which the pathways are being altered requires further investigation.The striking disruption in the wing formation drove us to further investigate the wing disc phenotypes driven by the *DVL1^1519^*^Δ*T*^ variant in comparison to control and wt*DVL1^1519^*^Δ*T*^ expressing wing discs.

### DVL1 variant disrupt morphology in imaginal wing discs

Based on the phenotypes we saw in the adult tissue that was expressing the *DVL1* variant, we were curious to look at tissue morphology at earlier stages. We saw evidence of tissue malformation in imaginal wing discs with three different Gal4 drivers (*dpp-, ap-, hh-Gal4*). Common abnormalities in wing disc morphology included apparent overgrowth and disrupted folding. The *hh>DVL1^1519^*^Δ*T*^ discs exhibited extra folds in both apicobasal and dorsoventral axes, showing a more complex and disorganized morphology in maximum projection and orthogonal views (Fig. 2, Fig. S2). In *ap>DVL1^1519^*^Δ*T*^ discs, along with the abnormal folds, the posterior side bent the discs basally (Fig. 2, Fig. S3). As the wing imaginal disc grows, its flat epithelial structure transforms, and folds begin to form at specific locations. The placement and development of these folds are regulated by various signaling pathways, including Wg and Jak-Stat (Villa-Cuesta *et al*., 2007; Sui *et al*., 2012; Johnstone *et al*., 2013; Wang *et al*., 2016; Sui and Dahmann, 2020). Fold formation also involves several cellular mechanisms, such as apical-basal cell shortening, microtubule redistribution, and localized degradation of the extracellular matrix (Sui *et al*., 2012; Wang *et al*., 2016). Additionally, computational studies have highlighted the role of differential growth between regions of the wing disc in fold formation (Tozluollu *et al*., 2019). In addition, the nuclei on the basal side appeared more punctate and spread apart, which could be indicative of cell death or migration (Dove, 2003; Bell and Lammerding, 2016). There was also a shift in the nuclear staining trending apically.

The basally directed folding of the posterior side of wing discs was very striking and consistent across wing discs. One possibility for this folding could be tension induced from half the wing discs being overgrown while the other half was normal. Therefore, we investigated the cell proliferation and cell death in the wing imaginal discs.

### Robinow Syndrome variant induce cell death

We observed changes in tissue morphology and investigated if altered proliferation or cell death contribute to the effects. To our surprise, there was no difference in cell proliferation as measured by pH3 staining across all genotypes tested. In contrast, we found induction of cell death after expression of either wt*DVL1* or *DVL1^1519^*^Δ*T*^. Quantification showed that the variant significantly increased cell death relative to control and wt*DVL1* (Fig. 5, Fig. S4, Fig. S5, Fig. S6). These results suggest that altered cell death could contribute to the apparent misfolding of the disc epithelium and the abnormal basal bending by changing tension within the disc as cells die. Furthermore, dying cells often extrude basally from tissues (Fadul and Rosenblatt, 2018), which we see in the cross sections of flip out expression clones (Fig. 4). These cells may contribute to the floating cells seen within the pupal blisters (Fig. 6).

### DVL1 variants disrupt dorso-ventral cell adhesion in pupal wings

The blistering in the pupal wings where DVL1 variants were expressed was intriguing. Previous studies have shown that the overexpression of Matrix metalloproteinase-2 (Mmp2) in the wing tissue can induce blisters (Sun *et al*., 2021). In that study, researchers hypothesized that for the dorsal and ventral layers of the wing to adhere, the basement membranes would need to be slowly degraded. If they were degraded too fast or not sufficiently, wing blisters would be induced. We had previously shown that Mmp1 was induced in imaginal wing discs (Gignac *et al*., 2023) and so we tested if this overexpression was sustained in the pupal wing and found that indeed the blisters were filled with secreted Mmp1. While Mmp1 preferentially cleaves the extracellular connections between adjacent cells and Mmp2 preferentially cleaves the connections between cells and the extracellular matrix, Drosophila studies have shown there is functional redundancy between them (Jia *et al*., 2014).

In this study we showed that basement membrane degradation is altered by the *DVL1^1519^*^Δ*T*^ variant (Fig. 8). We used Vkg-GFP protein trap as a proxy for the basement membrane, as *Drosophila* Collagen IV (Vkg) is the most abundant component. Vkg is present in the distal lumen of the L3 longitudinal veins during early pupal wing development (Murray *et al*., 1995), but is later eliminated by high Mmp1 activity (De Las Heras *et al*., 2018) as we also showed in our controls for this experiment (Fig. 8A-A’). What we see with the expression of *DVL1^1519^*^Δ*T*^ variant in the region between the L3 and L4 longitudinal veins of pupal wings suggests that the variant disrupts the regulation of Vkg and Mmp1 proteins during dorsoventral adhesion.

In our previous work we showed abnormal creases seen in adult wings that may be residual effects of the pupal blisters after they resolve (Gignac *et al*., 2023). This crease could be considered a result of wound healing and reepithelialization driven by the increased JNK signaling activity and Mmp1 secretion which would promote extracellular membrane assembly after wound formation (Holmbeck *et al*., 1999; Rämet *et al*., 2002; Galko and Krasnow, 2004; Srivastava *et al*., 2007). Disruptions in the regulation of wound healing mechanism driven by Mmp1 mutants showed that the wound healing is delayed and the reepithelialization was disrupted in these conditions, which also showed decrease in the deposition of Vkg-GFP to the basement membrane in the wound site (Stevens and Page-McCaw, 2012). We previously showed an increase in the JNK signaling readouts due to the expression of *DVL1^1519^*^Δ*T*^ variant (Gignac *et al*., 2023), which we showed to be dependent on the activity of JNK signalling in this study (Fig. 6). Along with this increase, our data on pre-expansion pupal wings that shows a high abundance of Vkg-GFP in the presence of *DVL1^1519^*^Δ*T*^ can be explained by the Mmp1 secretion into the blistered “wound” site, which would cause an increase in the deposition of Vkg-GFP. Mmp1 typically leads to the degradation of the basement membrane up until 3-6h APF, then gets inhibited (De Las Heras *et al*., 2018). The presence of ectopic Mmp1 puncta in wing discs expressing *DVL1^1519^*^Δ*T*^, along with the Mmp1-filled pupal wing blister, suggests that Mmp1 protein levels in the developing wing tissue are misregulated throughout wing development. Future investigations will look into the integrity of the pupal wing basement membrane in earlier pupal stages where wt*DVL1* or *DVL1^1519^*^Δ*T*^ variant are expressed to see if the overexpression of Mmp1, induced by the variant expression, leads to the altered degradation of the basement membrane and thus interferes with dorso-ventral cell adhesion in later pupal stages.

In summary, our work has shown that *DVL1* mutations implicated in RS can be modeled in *Drosophila* appendages where we can gain deeper understanding of the cellular consequences of dominant variant expression. Our model enabled assessment of pathological severity with the phenotypes we have shown. The defects in leg segments are particularly intriguing given that RS is characterized by shortened limb bones. Future work will delve into signaling pathways that control leg specification, growth and morphology.

## Materials and methods

### Drosophila Husbandry

Flies were raised on standard media. Stocks were kept at room temperature (∼22°C) and crosses were performed at 25°C or 29°C as indicated. Three Gal4 driver fly lines were used to induce transgene expression: *dpp-Gal4, UAS-GFP* /TM6B (Swarup, Pradhan-Sundd and Verheyen, 2015), *Dll-lacZ/Cyo; Hh-Gal4/TM6B* (Hall *et al*., 2017)*, ap-Gal4, UAS-GFP; +/SM6a∼TM6B* (generated for this paper by Katja MacCharles). Additional stocks used were: *;Vkg-GFP;* (Flytrap) (Blaquiere *et al*., 2018), *puc-LacZ* (Martin-Blanco *et al*., 1998)*, w^1118^* (RRID:BDSC_5905), *UAS-GFP*, *UAS-myr-RFP* (RRID:BDSC_7118)*, UAS-bskDN* (RRID:BDSC_6409) (Adachi-Yamada *et al*., 1999).

### Generation of somatic flip-out clones

To generate heat-shock inducible actin flip-out clones, *yw,hsFlp[122];;act>CD2>Gal4, UAS-RFP/TM6B* was used. A 10 min heat shock (37°C) was applied to larvae grown for two days after egg laying (25°C). After heat shock, the larvae were grown at 29°C and dissected when they reached third instar larval stage.

### Dissections

#### Wing imaginal discs

Third instar larval wing imaginal discs were dissected out of larvae in cold PBS. The top half of the larvae were separated and flipped inside out to expose the imaginal discs. Fat tissue and the guts were removed while leaving the imaginal tissue attached to the cuticle.

#### Pupal wings

Pupal wings were dissected as described by Bolatto and coworkers (Bolatto et al., 2017) with some modifications. 0hr APF pupae were collected and placed ventral side down onto a microscope slide covered with double-sided tape. The microscope slides were placed at 25°C until the desired stage of development. Pupal cases were opened with forceps and naked pupae were transferred onto a new microscope slide with double-sided tape and covered with 4% paraformaldehyde (PFA) in PBS. After a 30-minute incubation, pupal wings were dissected out and placed into a 4% PFA filled well of an HLA plate for an additional 15-minute incubation. All immunofluorescence experiments with pupal wings were performed in Greiner HLA Terasaki multiwell (54-well) plates rather than Eppendorf tubes.

#### Pharate Pupae

Pupae were gently pulled off the sides if vials and placed on double sided tape on a microscope slides. Forceps were used to pry open the operculum and tear away the case. Pharates were then photographed using a Leica dissection microscope and an iPhone.

#### Adult wings

Adult flies of the target genotype were collected in small glass vials filled with 70% ethanol. The wings were dissected using forceps in 95% ethanol in dissection dishes and mounted immediately. Aquatex (EMD Chemicals) was used as mounting media. The slides were baked overnight at 65°C incubator with small weights on them.

### Immunofluorescence staining, microscopy, and image processing

Tissue was dissected in phosphate-buffered saline (PBS) and fixed in 4% PFA at room temperature for 15 minutes. Samples were washed twice for 10 minutes with PBS with 0.1% Triton X-100 (PBS-T). Following a 1-hour block with 5% bovine serum albumin diluted in PBST at room temperature, samples were incubated overnight with primary antibodies at 4°C. The following primary antibodies were used:

### List of primary antibodies used in this study

| **Antibody** | **Dilution** | **Clone** | **Distributor** |
| --- | --- | --- | --- |
| Mouse anti-FLAG | 1:500 | M2 | Sigma |
| Rabbit anti-FLAG | 1:500 | SIG1-25 | Sigma |
| Rabbit anti-Dcp1 | 1:500 | Asp215 | Cell Signaling |
| Rabbit anti-PH3 Ser10 | 1:500 | D2C8 | Cell Signaling |
| Mouse anti-Armadillo | 1:50 | N2 7A1 | DSHB |
| Mouse anti-β-Galactosidase | 1:50 | 40-1a | DSHB |
| Rabbit-anti-GFP | 1:500 | A-11122 | Thermo Fisher |
| Mouse anti-Mmp1 | 1:100 (each) | 3A6B4, 3B8D12, 5H7B11 | DSHB |
| rat anti-DE-Cadherin | 1:50 | DCAD2 | DSHB |

Samples were washed twice for 10 minutes with PBS-T (0.5% Triton-X100 in PBS) and incubated with Cy3 and/or Alexa Fluor 647-conjugated secondary antibody (1:500, Jackson ImmunoResearch Laboratories), DAPI (4’,6-Diamidino-2-Phenylindole) and/or Rhodamine Phalloidin (Invitrogen™ R415). for 2 hr at room temperature. After two 10- minute washes, samples were mounted in 70% Glycerol in PBS or VECTASHIELD® Antifade Mounting Medium (Vector Labs) and imaged using a Nikon Air laser-scanning confocal microscope or a Zeiss LSM880 with Airyscan confocal microscope. Images were processed with ImageJ, FIJI software (Schindelin *et al*., 2012; Schneider, Rasband and Eliceiri, 2012) and are presented as Z-stack maximum intensity projections unless otherwise stated.

### Quantification of cell death and cell proliferation using Dcp1 and PH3 staining

After completing the staining and mounting steps, wing imaginal discs were imaged as described in the previous section. Using ImageJ software (Schindelin *et al*., 2012), the *ap-GFP* or *hh-Gal4* domains on wing imaginal discs were marked. PH3 or Dcp1 positive cells were counted within each region automatically using the Analyze → Analyze Particles tool after thresholding. The difference between the number of PH3 or Dcp1 puncta within the *apterous* or *hedgehog* domains were then plotted.

### Quantification of *puc-lacZ* signal intensity in wing discs

After completing the staining and mounting steps, wing imaginal discs were imaged as described previously. The signal intensity was measured by using the measurement tools in ImageJ software (Schindelin *et al*., 2012): Using a rectangular box of identical dimensions, mean β-Galactosidase signal intensity was quantified inside and outside (dorsal and ventral compartments) of the transgene expression domain. The ratio of signal inside/outside β- Galactosidase signal was used to determine *puc-lacZ* levels within the transgene expression domain.

### Quantification of Mmp1 puncta signal intensity in wing discs

After completing the staining and mounting steps, wing imaginal discs were imaged as described previously. Using ImageJ software (Schindelin *et al*., 2012), the *dpp-Gal4* domains of the wing discs were isolated in maximum projection images and the Mmp1 puncta was manually counted.

### Statistical analyses

Statistical analyses of the plotted data of mRNA and protein expression were done using one-way analysis of variance (ANOVA). For immunofluorescence experiments, paired (Arm stability within 1 wing disc) or unpaired (all other experiments) t-tests were performed to determine p-values. All statistical analyses were performed using GraphPad Prism 9.3.1 (GraphPad Software, San Diego, USA). Analyzed data with p<0.05 was considered statistically significant. Significance depicted as *p < 0.05, **p < 0.01, ***p < 0.001, ****p < 0.0001, ns = not significant.

## Acknowledgements

We are grateful to the Developmental Studies Hybridoma Bank (IA, USA) and the Bloomington Drosophila Stock Centers for providing antibodies and fly strains. The images in Fig. 1A-B and 2E were created with BioRender.

## Figure legends for Supplementary Figures

**Fig. S1 *DVL1^1519^*^Δ*T*^ induces abnormal morphology in male adult fly appendages and thorax**

(A-C) Male adults from crosses grown at 25°C of the indicated genotypes. Leg and wing defects are visible in the variant (C). Single third legs (T3) showing of the indicated genotypes. (A”-C”) Males from crosses grown at 29°C to drive higher transgene expression. Pharate pupae in C” was dissected from pupal case. (D) The length of T3 leg segments were measured and plotted for control, wt*DVL1* and *DVL1^1519^*^Δ*T*^ expressing flies, indicated by green, yellow and pink dots, respectively. Statistics were performed using Tukey’s multiple comparisons test. *p<0.05,

****p<0.0001. (E-G) Female and male adult body parts from crosses of the indicated genotypes grown at 29°C. (E-G) The male adult notum and scutellum. (E’-G’) Dorsal view of the whole male adult body. (E’’-G’’) Dorsal view of the whole female adult body.

**Fig. S2 *DVL1^1519^*^Δ*T*^ variant disrupt morphology in larval wing imaginal discs when expressed in the *hedgehog-Gal4* domain at 25°C**

(A-C) Wing imaginal discs from female larvae grown at 25°C and using *hh-Gal4* to express transgenes. Images showing DNA (DAPI, blue), F-actin (red) for (A, A’) control *UAS-RFP* (B-B’) wt*DVL1* and (C-C’) *DVL1^1519^*^Δ*T*^ wing discs mounted on bridged slides to retain their three-dimensional shape. (A-C) Maximum projections, (A’-C’) single plane images with orthogonal views of the tissue. Scale bar = 100 μm.

**Fig. S3 *DVL1^1519^*^Δ*T*^ variant disrupt morphology in larval wing imaginal discs when expressed in the *apterous-Gal4* domain at 25°C and 29°C**

(A-C) Merged confocal microscopy images showing the *ap-Gal4* UAS-GFP (green), UAS transgene expression domain (anti-FLAG, magenta), DNA (DAPI, blue), staining in (A) control,

(B) wt*DVL1* and (C) *DVL1^1519^*^Δ*T*^ wing discs grown at 25°C mounted on bridged slides to retain their three-dimensional shape. (A-C) Maximum projections, (A’-C’) single plane images with orthogonal views of the tissue. (D-F) Merged confocal microscopy images showing the *ap-Gal4* UAS-GFP (green), UAS transgene expression domain (anti-FLAG, magenta), DNA (DAPI, blue), staining in (D) control, (E) wt*DVL1* and (F) *DVL1^1519^*^Δ*T*^ wing discs grown at 25°C mounted on bridged slides to retain their three-dimensional shape. (D-F) Maximum projections, (D’-F’) single plane images with orthogonal views of the tissue. Scale bar = 100 μm.

**Fig. S4 *DVL1^1519^*^Δ*T*^ does not promote proliferation but causes elevated cell death when expressed in the *hedgehog-Gal4* domain at 29°C**

(A-C) Merged confocal microscopy images showing the *hh-Gal4* UAS transgene expression domain (anti-FLAG or RFP, red), DNA (DAPI, blue) and PH3 (white) staining in (A) control, (B) wt*DVL1* and (C) *DVL1^1519^*^Δ*T*^ wing discs grown at 29°C. (A’, B’, C’) The *hh-Gal4* domain of representative wing discs is shown with the RFP or FLAG staining images. (A’’, B”, C’’) PH3 staining for all genotypes are shown. (D) The number of PH3 puncta counted within the *hh-Gal4* domain are plotted. (E-G) Merged confocal microscopy images showing the *hh-Gal4* UAS transgene expression domain (anti-FLAG or RFP, red), DNA (DAPI, blue) and anti-Dcp1 (white) staining in (E) control, (F) wt*DVL1* and (G) *DVL1^1519^*^Δ*T*^ wing discs grown at 29°C. (H) The number of Dcp1 puncta counted within the *hh-Gal4* domain are plotted. Sample size is shown under each genotype. Statistics were performed with ANOVA. ****p<0.0001. Scale bar = 100 μm.

**Fig. S5 *DVL1^1519^*^Δ*T*^ does not promote proliferation but causes elevated cell death when expressed in the *apterous-Gal4* domain at 25°C**

(A-C) Merged confocal microscopy images showing the *ap-Gal4* UAS transgene expression domain (GFP), DNA (DAPI, blue), filamentous actin (F-actin, Rhodamine-Phalloidin, red) and PH3 (white) staining in (A) control, (B) wt*DVL1* and (C) *DVL1^1519^*^Δ*T*^ wing discs grown at 25°C. (A’, B’, C’) The *ap-Gal4* domain of representative wing discs is shown with the GFP. (A’’, B”, C’’) PH3 staining for all genotypes are shown. (D) The number of PH3 puncta counted within the *ap- Gal4* domain are plotted. (E-G) Merged confocal microscopy images showing the *ap-Gal4* UAS transgene expression domain (GFP), DNA (DAPI, blue), filamentous actin (F-actin, Rhodamine-Phalloidin, red) and anti-Dcp1 (white) staining in (E) control, (F) wt*DVL1* and (G) *DVL1^1519^*^Δ*T*^ wing discs grown at 25°C. (H) The number of Dcp1 puncta counted within the *hh-Gal4* domain are plotted. Sample size is shown under each genotype. Statistics were performed with ANOVA.

****p<0.0001. Scale bar = 100 μm.

**Fig. S6 *DVL1^1519^*^Δ*T*^ does not promote proliferation but causes elevated cell death when expressed in the *apterous-Gal4* domain at 29°C**

(A-C) Merged confocal microscopy images showing the *ap-Gal4* UAS transgene expression domain (GFP), DNA (DAPI, blue), filamentous actin (F-actin, Rhodamine-Phalloidin, red) and PH3 (white) staining in (A) control, (B) wt*DVL1* and (C) *DVL1^1519^*^Δ*T*^ wing discs grown at 29°C. (A’, B’, C’) The *ap-Gal4* domain of representative wing discs is shown with the GFP. (A’’, B”, C’’) PH3 staining for all genotypes are shown. (D) The number of PH3 puncta counted within the *ap- Gal4* domain are plotted. (E-G) Merged confocal microscopy images showing the *ap-Gal4* UAS transgene expression domain (GFP), DNA (DAPI, blue), filamentous actin (F-actin, Rhodamine-Phalloidin, red) and anti-Dcp1 (white) staining in (E) control, (F) wt*DVL1* and (G) *DVL1^1519^*^Δ*T*^ wing discs grown at 29°C. (H) The number of Dcp1 puncta counted within the *hh-Gal4* domain are plotted. Sample size is shown under each genotype. Statistics were performed with ANOVA.

****p<0.0001. Scale bar = 100 μm.

