## Supplemental figures for "Robinow Syndrome *DVL1* variants disrupt morphogenesis and appendage formation in a Drosophila disease model"

Fig. S1 ***DVL1*$^{1519ΔT}$ induces abnormal morphology in male adult fly appendages and thorax**

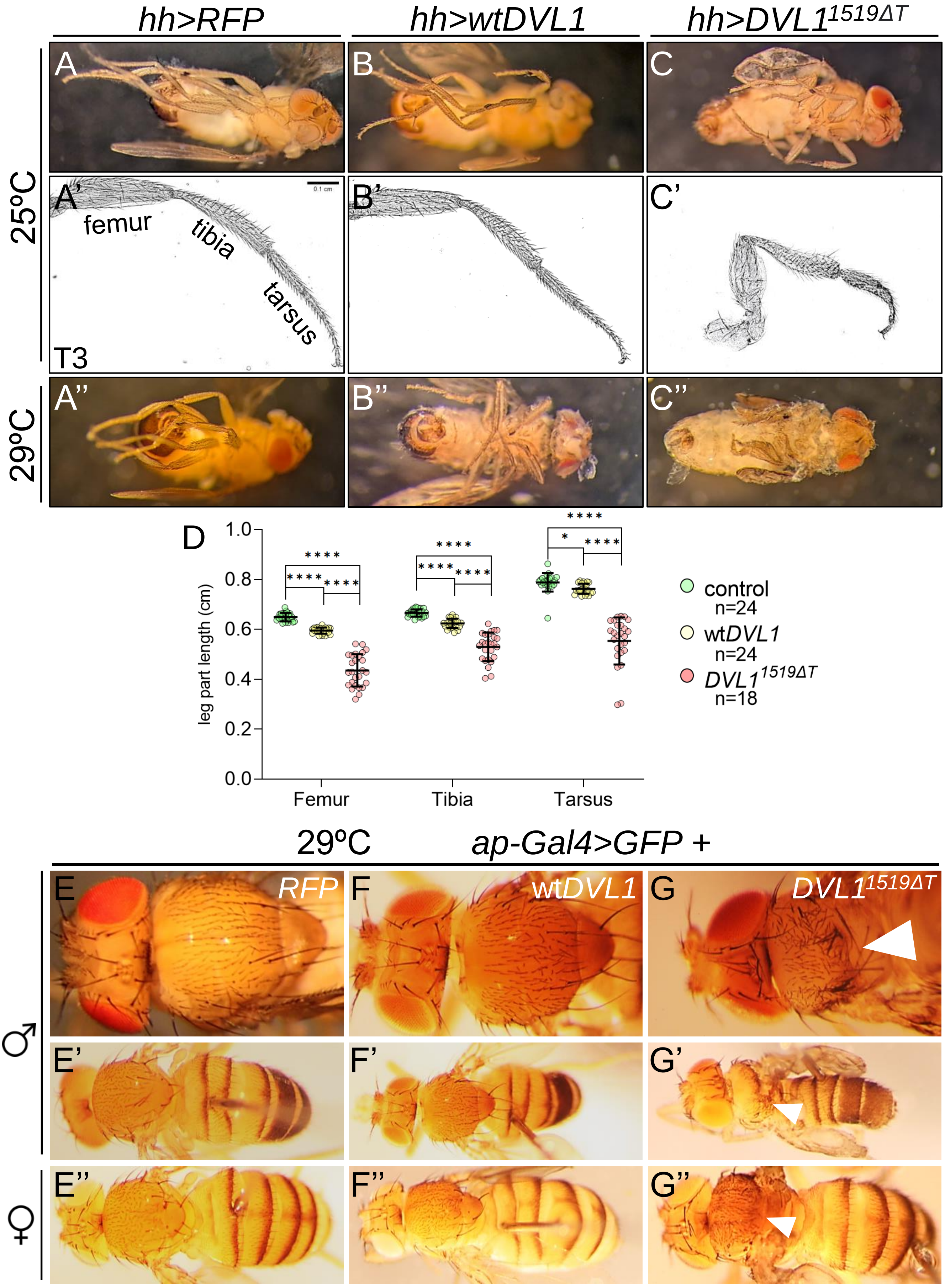

Fig. S2 ***DVL1$^{1519\Delta T}$* variant disrupt morphology in larval wing imaginal discs**

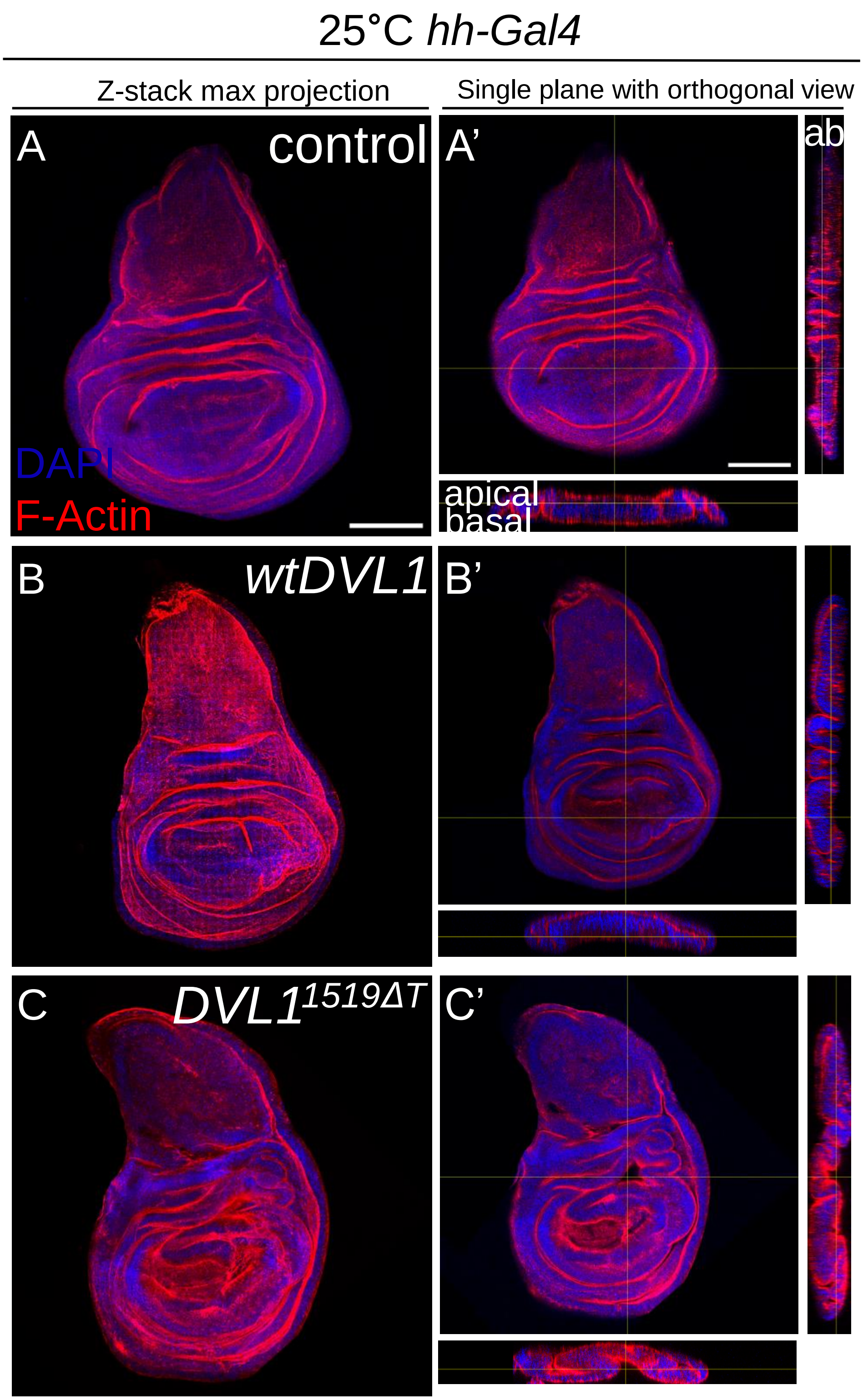

Fig. S3 **_DVL1_$^{1519\Delta T}$ variant disrupt morphology in larval wing imaginal discs when expressed in the _apterous-Gal4_ domain at 25°C and 29°C**

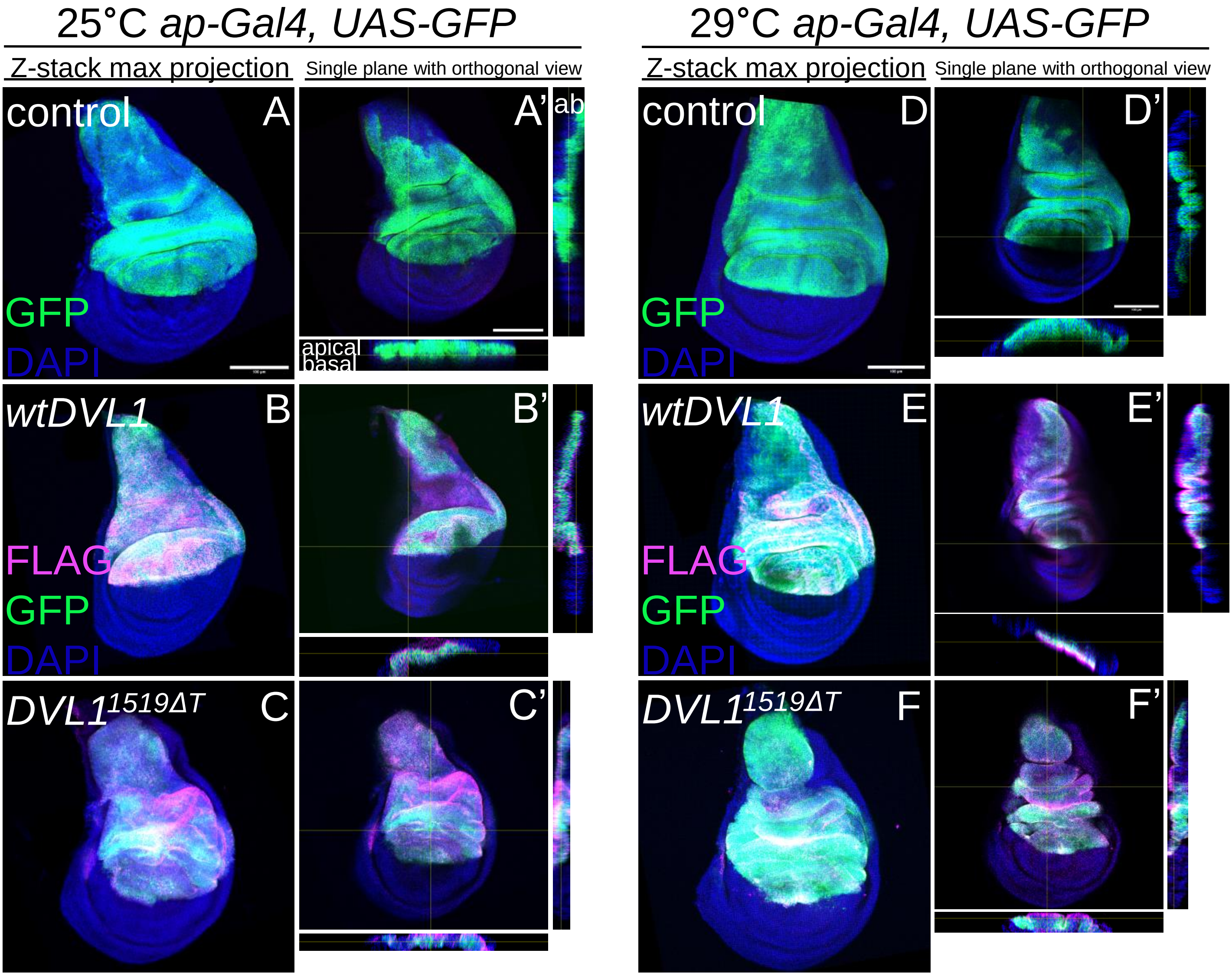

Fig. S4 ***DVL1$^{1519\Delta T}$* does not promote proliferation but causes elevated cell death when expressed in the *hedgehog-Gal4* domain at 29°C**

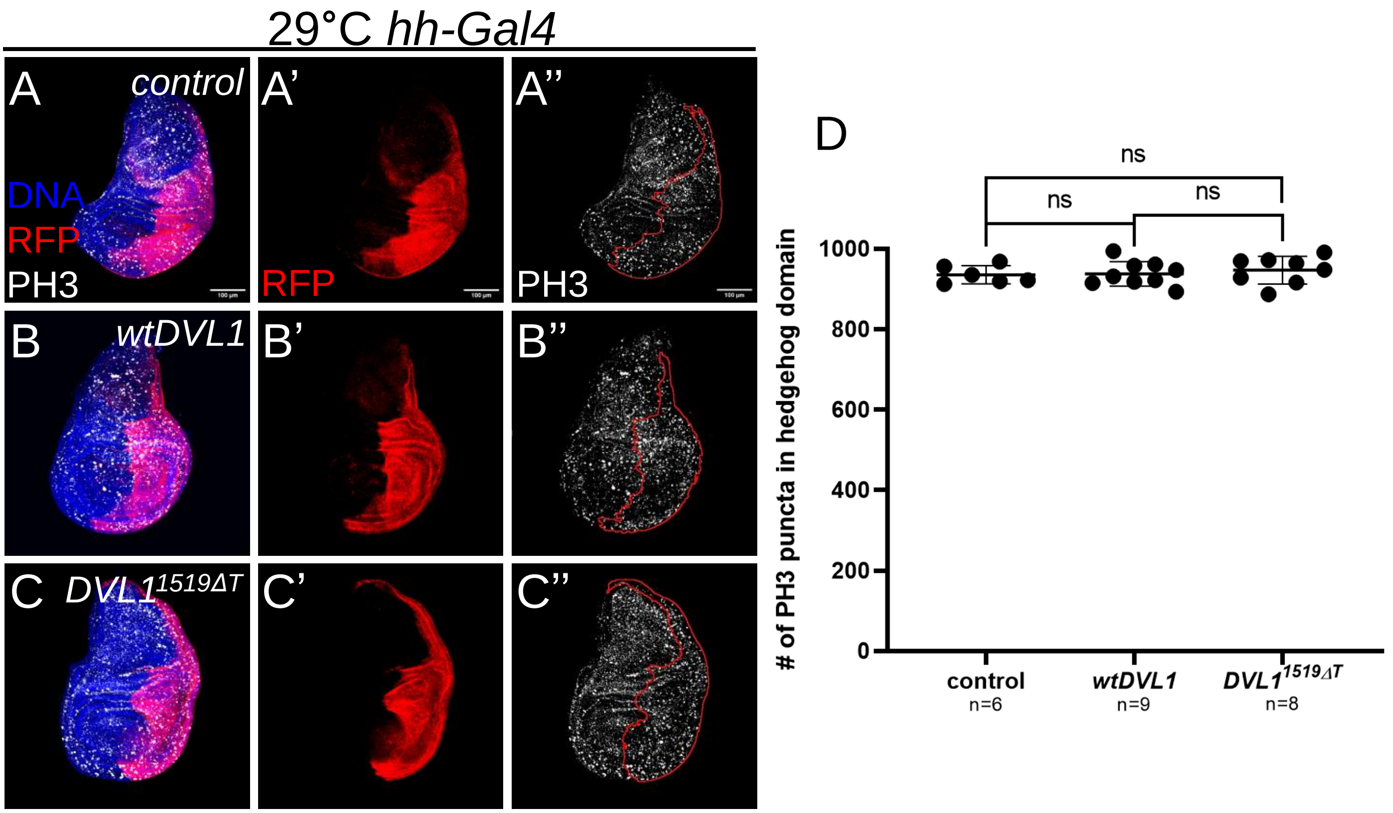


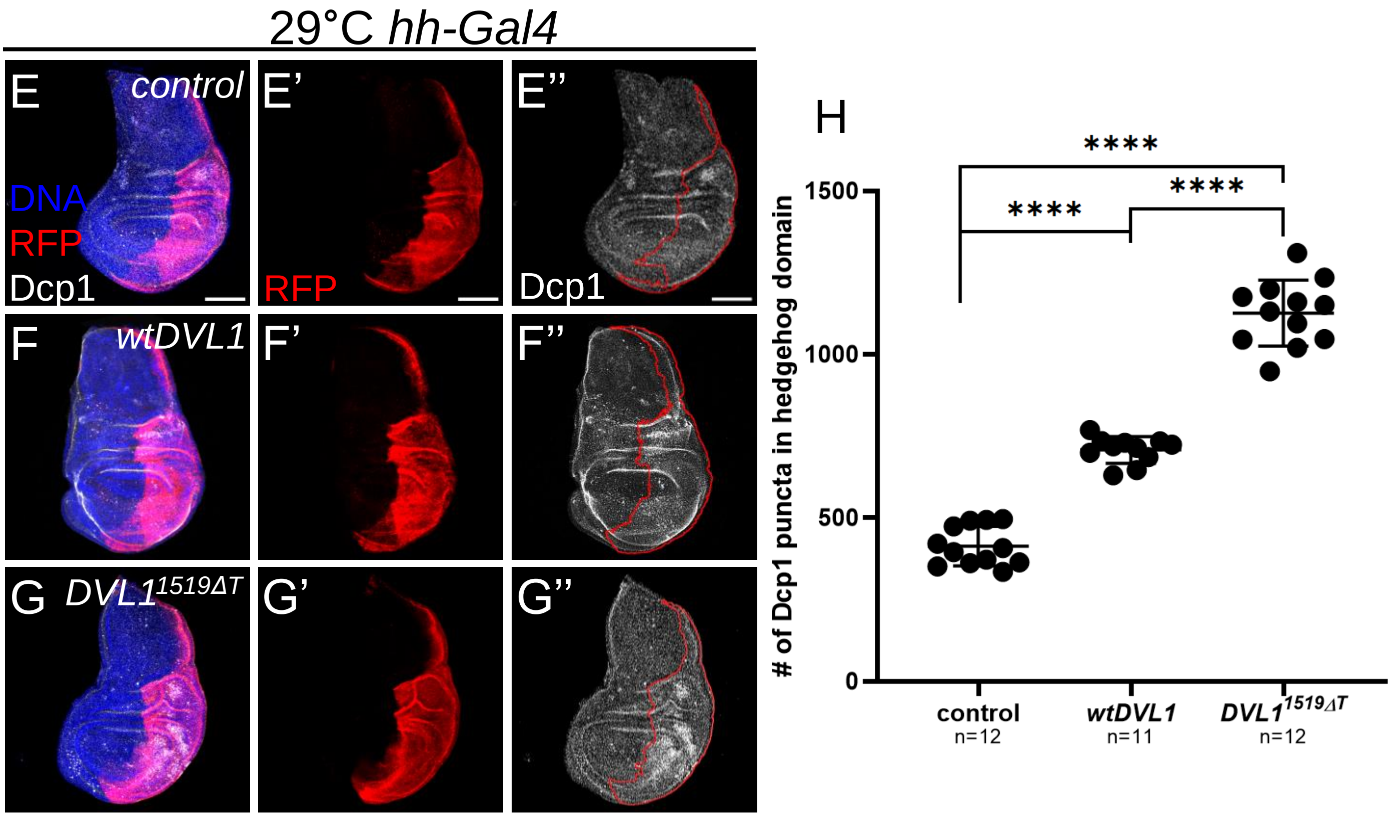

Fig. S5 ***DVL1$^{1519\Delta T}$* does not promote proliferation but causes elevated cell death when expressed in the *apterous-Gal4* domain at 25°C**

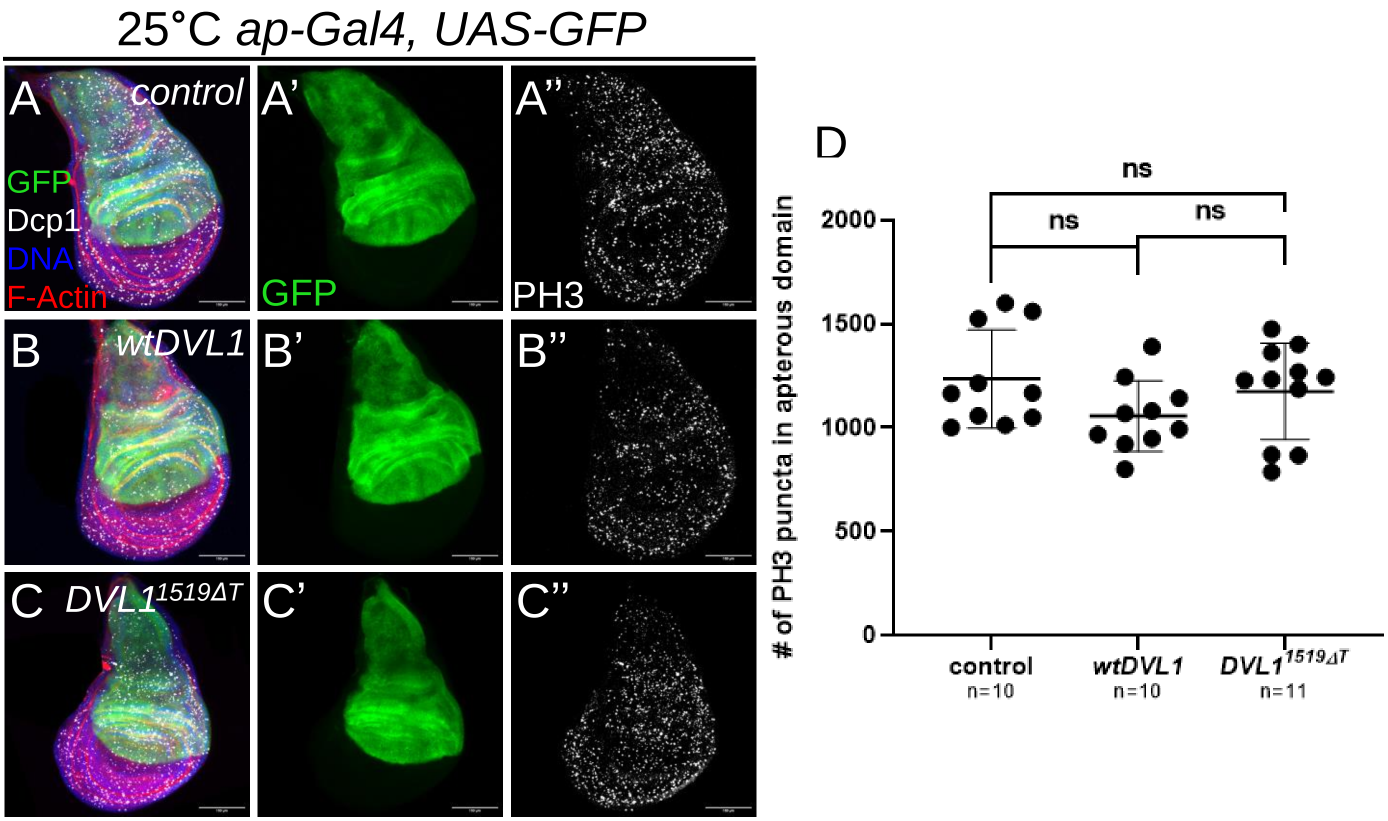


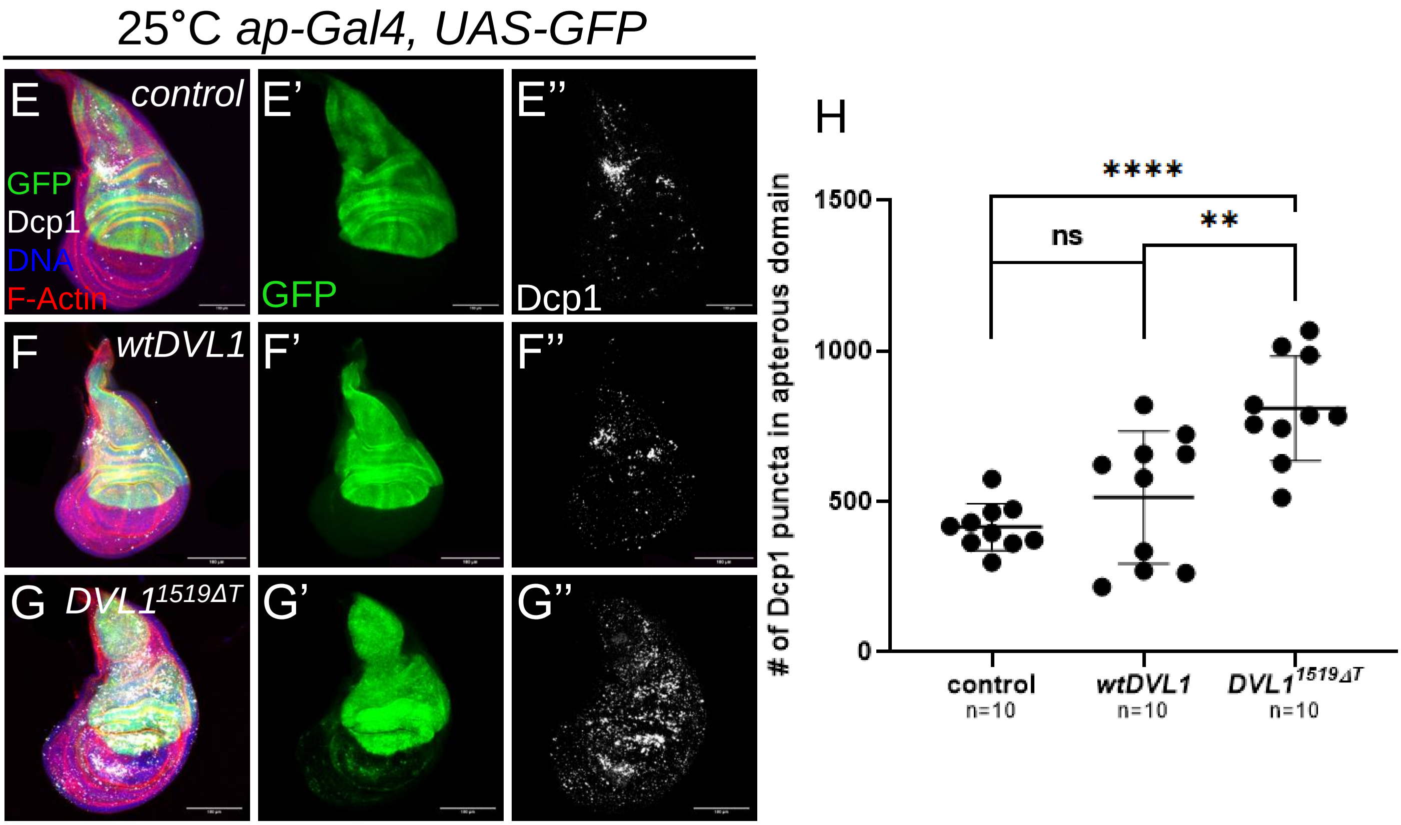

Fig. S6 ***DVL1$^{1519\Delta T}$* does not promote proliferation but causes elevated cell death when expressed in the *apterous-Gal4* domain at 29°C**

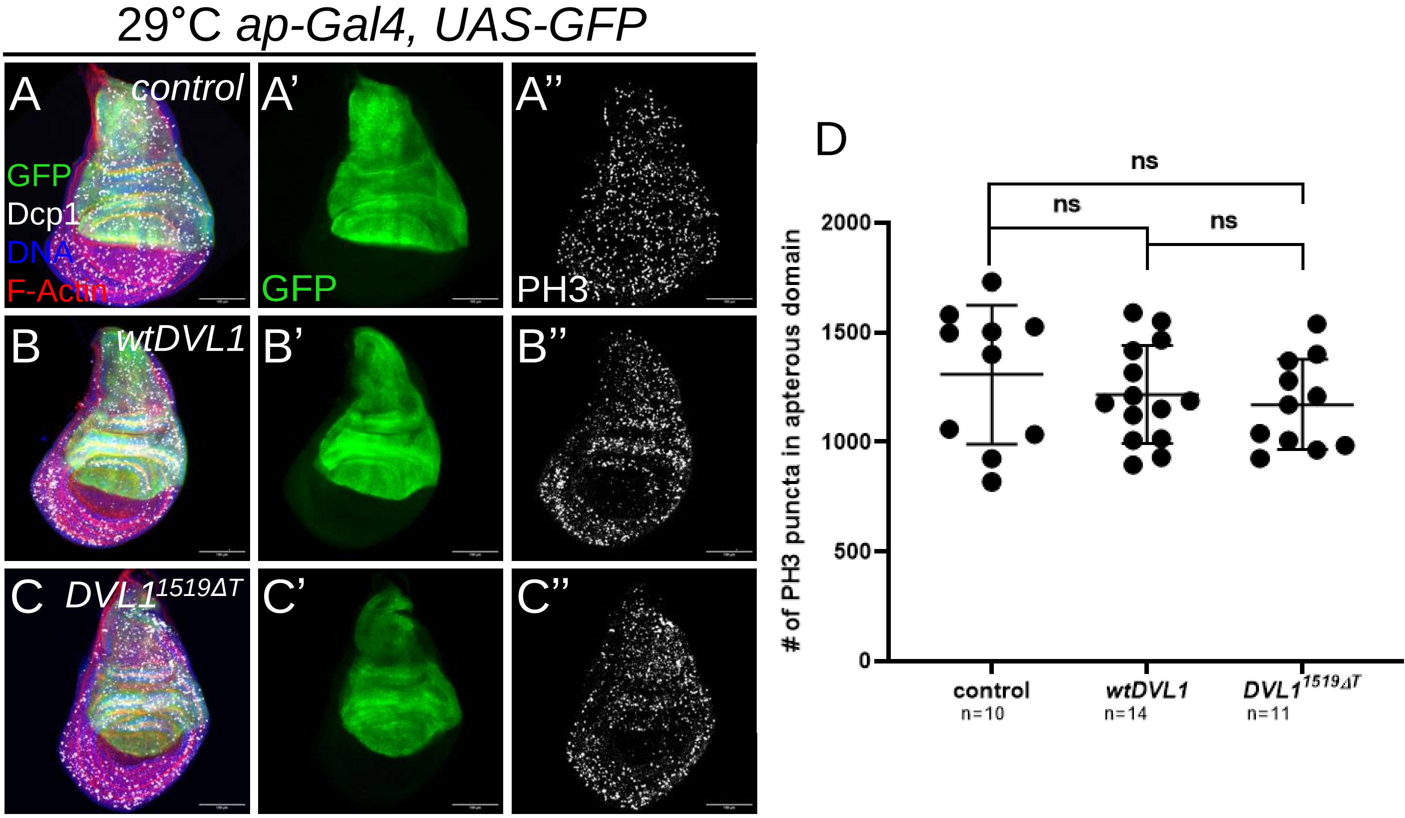


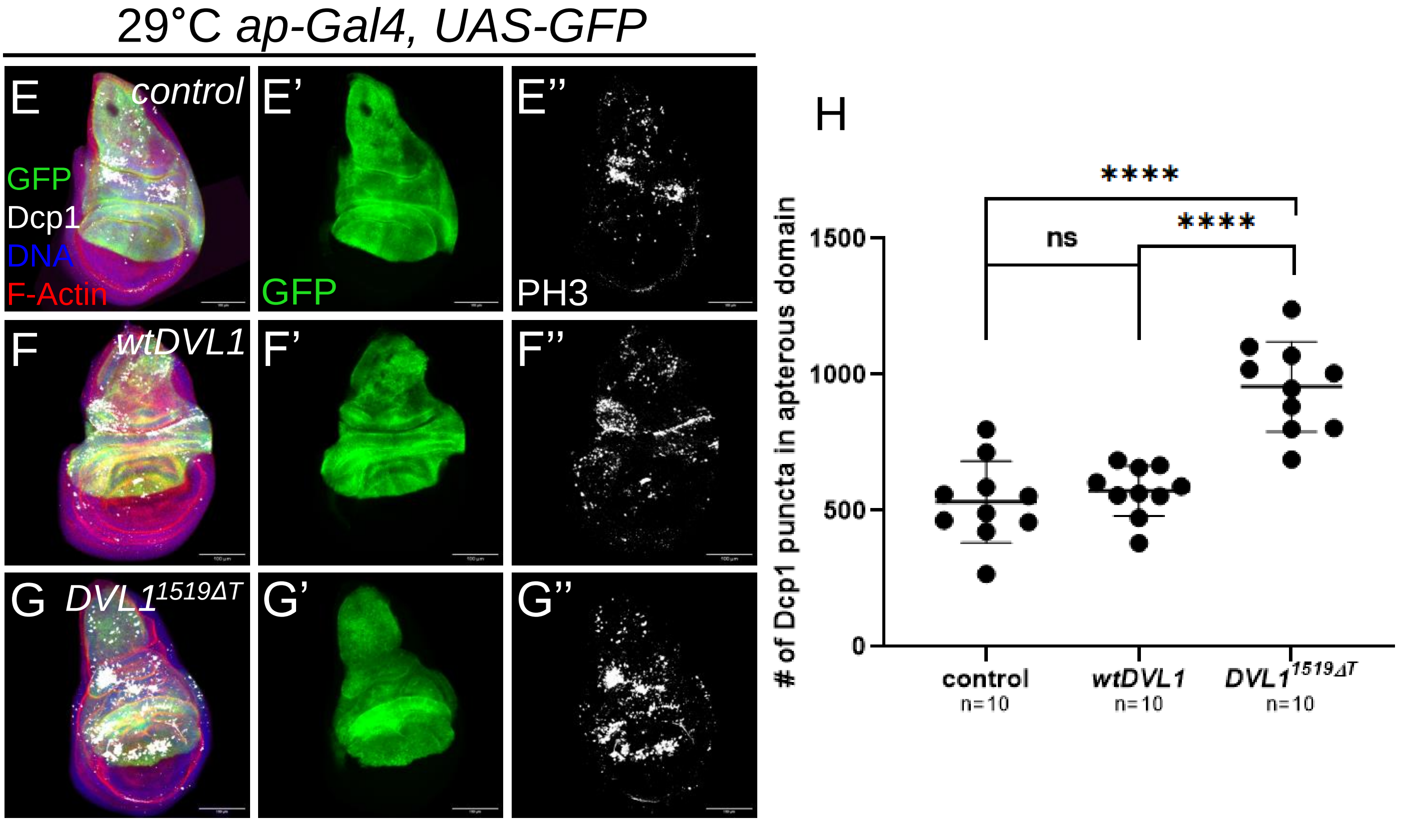
